# Visualizing and Quantifying microRNA Induced DNA Origami Separation at the Nanoscale

**DOI:** 10.1101/2025.08.15.670511

**Authors:** Chalmers C. C. Chau, Varun Gupta, George R. Heath, Christoph Wälti, Paolo Actis

## Abstract

Circulating microRNAs (miRNAs) are promising biomarkers for disease diagnosis, but their small size and instability hinder direct detection. The detection of small RNA, such as miRNA, using solid-state nanopores typically involves the binding of miRNA to a larger carrier molecule to generate detectable signals. However, this approach is prone to RNAse degradation in the environment leading to accidental digestion of miRNA prior to analysis. Here, we present an alternative approach based on DNA origami disassembly driven by toehold-mediated strand displacement (TMSD). Specifically, we designed a symmetric DNA origami dimer that undergoes TMSD-driven separation into monomers using miRNAs as invading strands. We visualized the real-time dynamics of dimer separation at high resolution using high-speed atomic force microscopy (HS-AFM), directly capturing nanoscale mechanical dynamics of the TMSD process that are inaccessible to ensemble or fluorescence-based measurements. Following the TMSD, single molecule nanopore sensing enables quantitative endpoint analysis of dimer separation by measuring the ratio of dimers to monomers. This direct read-out method enabled the multiplexed detection of miRNAs. Due to the near irreversible process of the TMSD, we performed the detection of miRNA in crude RNA tissue extracts under the presence of RNAse and showed that our approach allowed the robust detection of small RNA that is unaffected by the complex degrading environment.

## Introduction

The programmability of base pairing between nucleic acids has driven the advancement of DNA nanotechnology, enabling the construction of DNA nanostructures with precise sizes and shapes^1^. One notable example is DNA origami, which uses hundreds of short DNA oligonucleotide staples combined with a single-stranded DNA (ssDNA) scaffold to rationally design nanoscale structures^2–5^. Each DNA origami unit can be further assembled into higher-order supramolecular structures^6,7^ through blunt-end stacking^8^ or DNA strand hybridization^9–18^.

The connection between DNA origami units can be severed through methods such as toehold mediated strand displacement (TMSD)^10,19^. TMSD is an enzyme-free oligonucleotide strand exchange process where the starting hybridised dsDNA is composed of the original strand and a protector strand. The original strand includes an overhang region, or toehold, which is complementary to a third, invading strand. This invading strand is complementary to the original strand. The process initiates when the invading strand partially hybridises to the toehold and, through branch migration, displaces the protector strand, such that the resulting dsDNA no longer contains the toehold^20,21^.

The programmability of DNA origami also enables the formation of DNA:RNA heteroduplexes, which, when paired with an appropriate readout strategy, can be used for the specific detection of RNA molecules, including microRNAs (miRNAs)^14,22–27^. The miRNAs are a class of small, non-coding RNAs with average length of 19-25 nucleotides ^28,29^. They are highly conserved and play an important role in post-transcriptional gene regulation through translational repression and mRNA degradation^30^. As the expression profile of miRNA is intimately related to the cell status^31^, and therefore circulating miRNAs are clinical biomarkers for diseases including infectious diseases, cancers and physical injuries like traumatic brain injury (TBI)^32–35^. Direct detection of miRNA requires sensitive approaches, as they are present at sub-nanomolar concentrations in bodily fluids and are prone to degradation by ribonucleases (RNases) ^36,37^. They are typically quantified by hybridization-based microarrays and RT-qPCR^38^. More recently, innovative approaches such as nanopore sequencing have been successfully employed for miRNA detection^39^.

Nanopore sensing is a label-free biophysical technique allowing single molecule analysis^40–51^ that can provide a quantitative readout^52^, and it has been successfully employed for the fingerprinting of DNA origami^16,53,54^. The detection of miRNA with nanopores relies on a functional carrier such as magnetic bead, gold nanoparticle, DNA or DNA origami^45,46,55–58^. The carriers change size or conformation upon miRNA binding, and these changes can be “read” using a nanopore sensor. However, these approaches are susceptible to RNase-mediated degradation of the bound miRNA. While the addition of target miRNA to the nanostructures in an RNase-free environment and subsequent storage under clean conditions is stable, any RNase contamination, either during storage, sample handling or during measurement from equipment can degrade the hybridized miRNA and result in false negative read-outs.

Here, we propose a DNA origami strategy which is compatible with a nanopore readout and allows for the selective detection of miRNA in crude RNA extract. We designed and validated a DNA origami dimer formed through the dimerization of DNA origami tiles.^16–18^ The connection between the tiles consists of dsDNA linkers containing toeholds. Upon exposure of the dimer to the target oligos or miRNAs, the dimer is disassembled into two monomers. A nanopore sensor in combination with a polymer electrolyte^16,40–43,50,59^ can quantitatively monitor the relative concentration of the dimer and monomer tiles, and infer the miRNA presence and concentration. The polymer electrolyte method allowed us to detect the translocation of DNA origami in its native buffer without the need of high salt conditions, thus avoiding potential instability of the structure. We observed that both DNA and RNA can be used to separate the dimer, and miRNA can also be used when its secondary structure is disrupted. We also demonstrated that detection can be carried out under RNAse contamination at the point of measurement. The presence of RNAse does improve our detection sensitivity as the DNA origami are resistant to its action, and its presence eliminates background signal caused by small RNAs. In our approach, whether or not the miRNA is bound to the DNA origami after hybridisation does not affect the readout as long as the miRNA has already disassembled the dimer.

In addition to the nanopore readout, we directly visualised the TMSD reaction in real time using high speed-atomic force microscopy (HS-AFM)^60,61^. This enabled single molecule mapping of the stepwise separation of the dimer, revealing the dynamic progression of strand displacement and the positional preferences of linker regions. These observations provide unique mechanistic insights into the nanomechanics of TMSD at the DNA origami interface.

## Results and discussion

### DNA origami-based oligonucleotide sensor

A DNA origami dimer is generated from two DNA origami monomer tiles – the left and the right monomers (Figure 1a). The left and the right monomers are symmetrical tiles, and their design is based on previously published structures^16–18,59^. For dimerization, the staple sets of the edges of these monomers are modified with additional complementary ssDNA overhangs, such that a dimer can be formed through strand hybridisation^10–15^. The left and the right monomers are identical and composed of the same staples, except for the overhang edge’s staples; the resultant origami has a unique orientation reference marker, as shown in Figure 1.

**Figure 1.**
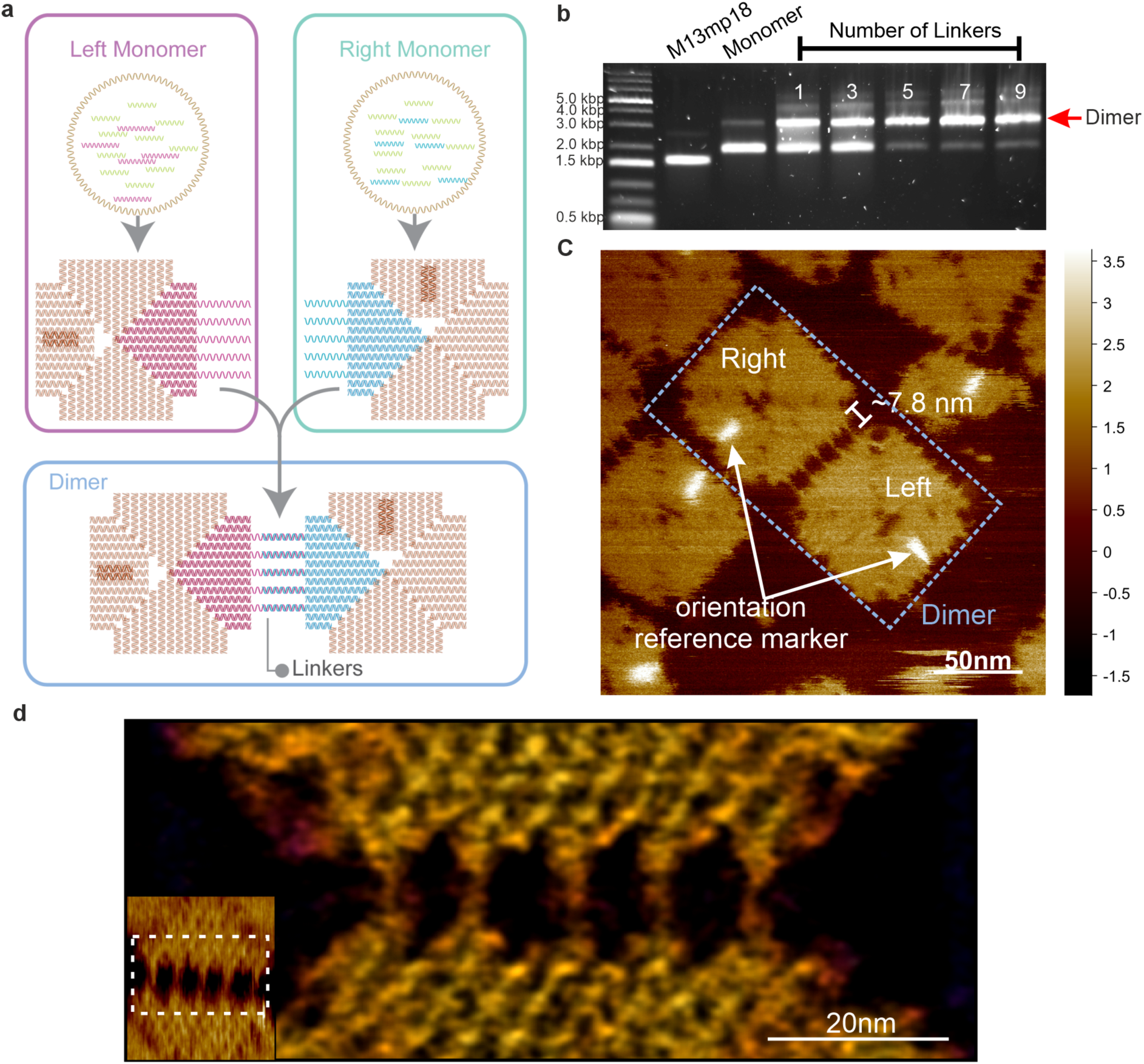
Design and characterisation of DNA origami dimers. (**a**) Schematic illustration on the formation of the 5-linker-dimer. The process involves two DNA origami monomers, the left (red) and the right (blue) DNA origami monomers, the edges of both left and right monomers contain five ssDNA overhangs that are 24 nt and 16 nt long respectively, these overhangs extend away from the main tile body of the origami. Additionally, the left monomer’s edges contain extra 8 nt at the 5’ end to act as toeholds after hybridisation. After the synthesis and purification of the monomers, the left and right monomers are mixed at equal portion and undergo a thermal incubation step to form the asymmetric 5-linker-dimer. (**b**) Agarose gel electrophoresis screening for the optimal number of linkers necessary to form dimers, 5 and more linkers led to high yield of dimers, as can be observed from the monomer bands that are a lot fainter than for 1 and 3 linkers. (**c**) AFM imaging of the resultant dimer. Through the unique orientation reference marker of the dimer, it allows us to confirm the formation of the dimer is via designated linkers. The length of the linkers is approximately 7.8 nm which is close to the theoretical length of 24 bp dsDNA (∼8.2 nm; 0.34 nm/bp). (**d**) Combining HS-AFM (inset) and LAFM approaches to visualise the five connecting linker strands.

In principle, the formation of the dimer can be carried out with a maximum of 11 linkers^16–18^. We screened the formation of dimers with 1, 3, 5, 7 and 9 linkers to identify the minimal number of linkers needed to generate a high yield of dimers (Figure S1a) and confirmed that dimer can be formed with 5, 7, and 9 linkers at high yield, while 1 and 3 linkers are not sufficient (Figure 1b, Figure S1b-c). Based on these results, we selected 5 linkers to generate dimers.

Atomic Force Microscopy (AFM) was used to characterise the formation of dimers. As can be seen in Figure 1c, the resultant dimers were asymmetric and connected with 5 linkers. Both the left and the right monomers could be identified, confirming the dimerization of the monomers occurred through the designated linkers^8^. We note that the left and the right monomers could self-dimerize to form dimeric structures through blunt end stacking^3,8,18^. However, these unintended stackings occurred predominately at the edges modified with overhang (Figure S2), and make up less than 10% of dimers. We also employed high-speed AFM (HS-AFM) to study the kinetics of dimer assembly and disassembly as well as localization-AFM (LAFM)^62,63^ to enhance the spatial resolution of the topographic features as shown in Figure 1 (d). Figure 1d shows the high-resolution localized visual of the DNA for all 5 linkers. On rare occasions, we observed that the DNA linkers crossed to form an “X” shaped connection between the 2^nd^ and 3^rd^ linker as shown by the LAFM at high-resolution (Figure S3a). While rare, it is not unexpected since all 5 linkers have the same sequences and are complementary. Additional high-resolution LAFM images of the origami can be found in Figure S3. The DNA linkers’ length was found to be 7.8 nm as shown by AFM and LAFM images, in good agreement with the theoretical length of 8.2 nm (0.34 nm/bp, 24 bp) per DNA linker.

The ssDNA overhangs of the left monomer contain additional nucleotides (8 nt) at the 5’ direction to act as toeholds (Figure 2a)^64–67^. After dimerization, the toeholds are generated through the hybridisation between the original strands (left monomers) and the protector strands (right monomers). The toehold in the linkers allows the dimer to be separated through toehold mediated strand displacement (TMSD) reactions^20,21^. The TMSD reaction follows a sequence of steps, as shown in Figure 2a. The speed of the strand displacement is dependent on toehold length, and for toeholds of 6 nt and above, the rate constant saturates^64–67^.

**Figure 2.**
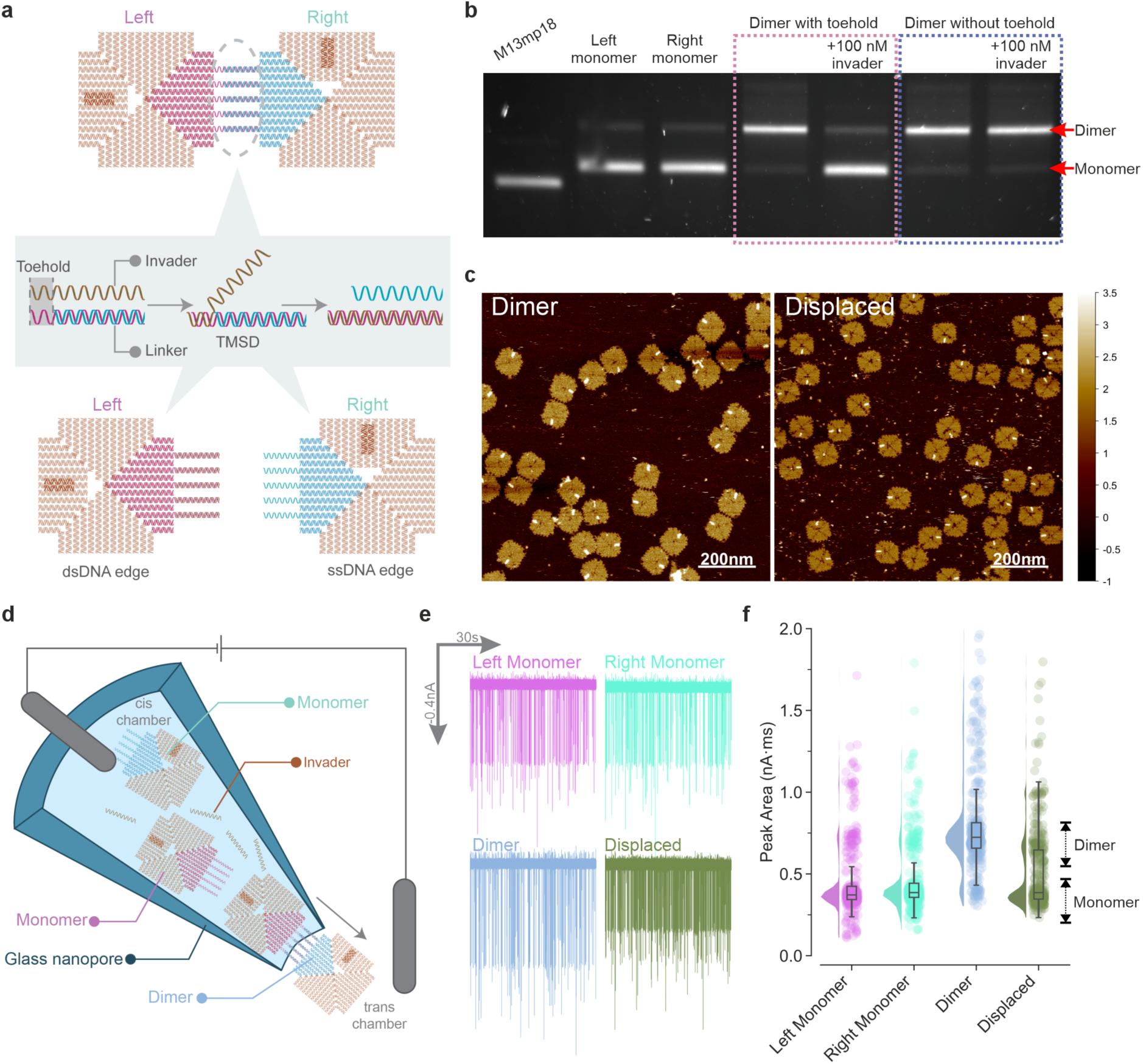
Characterising the TMSD separated DNA origami dimers. (**a**) Schematic illustration of the strand displacement mechanism through TMSD. The dimer is formed by hybridisation of the left and right monomers via five linkers, each linker contains an 8 nt toehold (ssDNA region) followed by fully hybridised dsDNA between the left and right monomers. The invader strand is fully complementary to the linker including the toehold. Upon addition, the invader strand initially hybridises to the toehold region, forming a three-stranded complex. Subsequent branch migration of the invader strand displaces the previously hybridised strand in the dsDNA region. This TMSD process occurs across all five linkers, leading to the dissociation of the dimer into two monomeric units. After displacement, the left monomer contains double-stranded DNA (dsDNA) overhangs, while the right monomer presents multiple single-stranded DNA (ssDNA) overhangs. (**b**) Gel electrophoresis analysis of the displacement of the dimer. The dimers were displaced by the introduction of invader at 100 nM, as evidenced by the significant increase and corresponding decrease in intensity of the monomer and dimer band, respectively. A dimer without toehold could not be separated by the invader at 100 nM, indicating that displacement was not spontaneous at room temperature. (**c**) AFM imaging of the dimer and the dimer after displacement. (**d**) The schematic illustration of the translocation of the monomers and dimer through nanopores. The DNA origamis were diluted in the origami buffer to 9.4 ng/µl (2 nM for dimer, 4 nM for monomers equivalent). The DNA origami was loaded into the cis chamber of the nanopore, a voltage of -500 mV was used to cause the translocation of DNA origami from the cis to trans chamber. The trans chamber is composed of the PEG polymer electrolyte. (**e**) The translocation trace of the left monomer (top left), right monomer (top right), dimer (bottom left) and the displaced dimer (bottom right). (**f**) The translocation event peak area comparison. The left and right monomers, and displaced dimer had a peak area median of approximately 0.4 nA·ms and the dimer had a peak area median of 0.7 nA·ms. The distribution was estimated by kernel density estimation.

The TMSD approach to separate the dimer was tested via mixing 2 nM of the dimer with 100 nM of invader strands (50× excess). The samples were incubated at 25°C for 30 mins followed by storage at 4°C until analysis. Gel electrophoresis and AFM were used to characterize the yield of the TMSD displacement reaction (Figure 2b-c, Figure S4). The dimers separated into monomers after the TMSD reactions, and where the toeholds were absent from the linkers, the addition of the invaders failed to separate the dimers.

Previously, we employed nanopore translocations to analyse the structures of DNA origami tiles, showing that the nanopore could differentiate and fingerprint supramolecular DNA origami structures^16^. Here, we utilised glass nanopores to quantify and differentiate monomers and dimers (Figure 2d). Our nanopore sensing approach uses a poly(ethylene) glycol (PEG) polymer electrolyte, and thus allows us to perform single molecule detection with high signal-to-noise ratio at physiological relevant salt condition^40–43^ (∼0.1 M monovalent salt). In contrast to many other nanopore based studies requiring monovalent salt concentration of 0.1-4 M^42,44,51,57,68^, the PEG electrolyte in the trans chamber allowed us to carry out the nanopore translocation experiments at low electrolyte concentrations in the cis chamber, without affecting the signal to noise ratio^40–43,50^ (the translocation buffer is only composed of: 10 mM Tris-Acetate, 15 mM Mg-Acetate, 1 mM EDTA, pH 8.0).

The nanopores used in this study had a diameter of approximately 50 nm, and their fabrication was highly reproducible, as confirmed by the voltammogram analysis (Figure S5a–b). No translocations were observed when the measurements were carried out in pure buffer (no DNA origami present) (Figure S5c). Following folding and purification, the DNA origami structures were diluted to 4 nM for monomers and 2 nM for dimers (approximately the same number of monomers in solution,). The samples were then translocated using a range of applied voltages, from –100 mV to –700 mV, to drive the negatively charged DNA origami from the cis to the trans chamber (Figure 2e, Figure S5d). A voltage bias of –500 mV was selected as the standard for further translocation experiments, as this voltage led to 300+ translocation events per minute, while minimizing the risk of nanopore clogging. The translocation of the dimer elicited a single molecule event with a larger current amplitude magnitude and slightly longer dwell time when compared with the monomer (Figure S5e). The most effective method for distinguishing between monomer and dimer events was by analysing the peak area of the translocation signals, consistent with previous findings from our group^16^. The dimer population exhibited a peak area of approximately 0.7 nA·ms, while the monomer population centred around 0.4 nA·ms (Figure 2f).

Both short ssDNA and RNA oligos can trigger a TMSD^65,69–72^ reaction by forming either a dsDNA duplex (denoted as ssDNA:ssDNA) or a RNA:DNA heteroduplex (denoted as RNA:ssDNA). We conducted a screening experiment to assess the displacement of the dimer using varying concentrations (2–25 nM) of ssDNA and RNA invaders (Figure S 6). The ssDNA and RNA oligos were identical in sequence, except for the substitution of thymine (T) with uracil (U) in the RNA. For both ssDNA and RNA invaders, the proportion of dimer remaining changed depending on the invader concentration (Figure S6c, e). Overall, the ssDNA invader exhibited higher displacement efficiency, leaving only about 50% of dimers remaining at a 6 nM concentration, compared to approximately 65% remaining for the RNA invader under the same concentration. In all tested concentrations of invaders, RNA invaders were less efficient than ssDNA at separating the dimers. At concentrations of 15 nM and above, both RNA and ssDNA invaders achieved complete dimer displacement.

We employed a fluorophore-quencher pair covalently attached to the linkers to characterise the kinetics of the TMSD. Upon displacement, the fluorophore was separated from the quencher, enabling real-time tracking of the reaction using a fluorescence plate reader over several hours (Figure S7). In agreement with a previous study^65^, the ssDNA:ssDNA reaction reached equilibrium slightly faster than the RNA:ssDNA reaction at all tested concentrations (5, 15, and 25 nM), as reflected by the rate constants: 2.45×10^5^, 1.41×10^5^ and 0.96×10^5^ M^-1^·s^-1^ for ssDNA invaders, compared to 1.15×10^5^, 0.87×10^5^ and 0.66×10^5^ M^-1^·s^-1^ for RNA invaders, respectively.

### Visualising TMSD

The separation of the dimer relies on TMSD^73–76^. The molecular details of this process are typically probed through fluorescent based bulk assays (Figure S7), single molecule fluorescence, force spectroscopy or computational simulation^77,78^. However, directly observing TMSD in action remains challenging due to the difficulty of imaging oligonucleotide dynamics with sufficient resolution and over relevant timescales. Here, leveraging the dimer origami’s design and the high spatial-temporal resolution of HS-AFM, we directly visualize the TMSD process that leads to the separation of the dimer. This allows us to capture its dynamics and reveal positional preferences of the linkers involved in displacement.

HS-AFM imaging of the dimer, connected by 5 DNA linkers (L1-L5), showed these linkers remained stable and connected during repeated AFM imaging for more than 840 frames (Figure S8-9, Supporting Movie 1-3). To monitor the TMSD process in real-time, DNA or RNA Invader strands were added *in situ* to the HS-AFM imaging buffer at a final concentration of 10 nM without interrupting imaging. Following the addition of the invader strands, the dimers’ linkers were observed to disconnect and separate from their neighbouring linkers at different time points. (Figure 3a-c, Figure S10-14, Supporting Movie 4-9).

**Figure 3.**
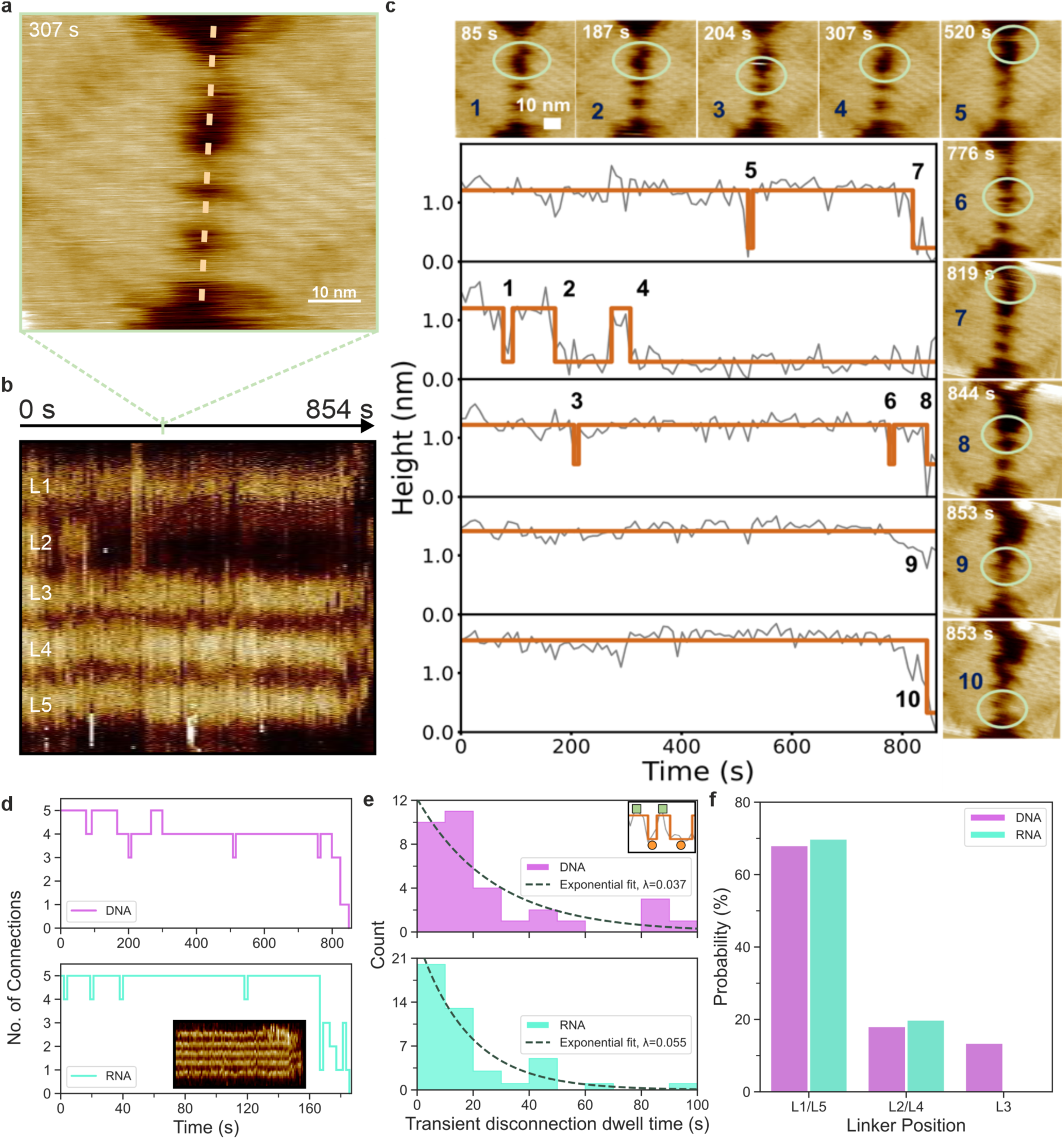
HS-AFM tracking of the separation of the dimer. To track the TMSD-based separation of the dimers, a single dimer was tracked in HS-AFM, and the separation was initiated by directly adding a final concentration of 10 nM of invader strands into the imaging buffer. (**a**) A snapshot of the dimer at 307 seconds after the initiation of the displacement. (**b**) Kymograph profile of HS-AFM video (Supporting Movie 4) to visualize the separation of each linker (L1, L2, L3, L4, L5) for the dimer. All 5 linkers were monitored over 854 seconds. (**c**) Kymograph height profiles along each individual linker over time with HS-AFM images selected at time points corresponding to on/off events (1-10). (d) Time evolution of the number of connections for origami separated by DNA invaders (top; analysis from panel c) and origami separated by RNA invaders (bottom; analysis from Supplementary Figure 12 and Supporting Movie 7). (e) Through HS-AFM, we observed the linkers changed from connected (green square; inset) to disconnected (orange circle; inset) then back to connected (green square; inset). This pattern of transient disconnections occurred multiple times before the irreversible separation of the dimer. Most transient disconnections (linker state from connect → disconnect → connect) lasted less than 30 seconds, with a higher reconnection rate with RNA invaders compared to DNA invaders. (**f**) The linker’s position influenced the probability of getting separated first (between L1-L5) during the TMSD process. The bar graph shows that the outer linkers (L1/L5) are more prone for disconnection after TMSD is initiated.

To measure the TMSD molecular process and dynamics, we generated kymographs^79^ from aligned HS-AFM movies to profile the topographic height of the linker regions over time (Figure 3b). This allows us to observe their separation after the invader strands were added to the imaging buffer (Figure 3a-c). The kymograph profile (Figure 3b) is obtained by measuring the height values along the dashed line in Figure 3a. In this kymograph, each vertical line corresponds to a HS-AFM frame at a specific time point. When the linkers are connected, the kymograph shows a higher height response at that location, whereas disconnection results in a decrease in the linker region height. The height profiles of all five linkers from the kymograph (Figure 3c) were analysed as binary states (0 = disconnected, 1 = connected), using a cut-off threshold of 0.75 nm in height. Figure 3c shows the 10 separation events that have been observed at different time points, as marked in the plot and inset images (1-10).

The first separation event was observed at around 75s (state-0), when the L2 linker became disconnected (Figure 3b, inset image 1). This linker re-established the connection within a few seconds (reverse back to state-1) and stays connected for approximately 20s. It is then disconnected again at around 100s (state-0). Similarly, other short, transient disconnection events were observed at different linker positions and at different time points (Figure 3c,d), before the dimer separated permanently through the disconnection of all L1-L5 linkers (t=853s). We also observed that reconnections could be formed between linker at different positions (e.g. between L1 with L3) as shown in Figure S11 (Supporting Movie 5). Importantly, these transient disconnections were not observed in the absence of invaders (Figure S8-9, Supporting Movie 1-3), strongly indicating that these transient events are due to the invaders’ interactions with the linkers. Figure 3d summarises the number of connected linkers over the HS-AFM tracking time for both DNA (Figure 3c, Supporting Movie 4) and RNA (Figure S12, Supporting Movie 7) invaders. In both cases, transient disconnections were observed, with the number of connections fluctuating between five and four for an extended period, followed by the rapid dissociation of the remaining linkers. We quantified the dwell times of the transient disconnections at different linker positions (Figure 3e). In both cases, for DNA and RNA invaders, the number of disconnections decreases exponentially with dwell time, suggesting a stochastic process with reconnection rates of 0.037 and 0.055, corresponding to half-lives of the disconnected states of around 18 s and 37 s, respectively.

The position of the first linker to separate (between L1-L5) after the addition of the invader strands were quantified, our data show that for both DNA and RNA invaders, the outer linkers (L1/L5) are more likely to separate first during the TMSD reaction (Figure 3f). Although, this could partly be an unintended artefact of HS-AFM imaging, when the tip moves from the flat mica surface onto the elevated outer linkers (L1/L5), the sudden change in topography may inject small amounts of mechanical energy to the exposed outer linkers, this effect is most pronounced when the fast-scan direction is perpendicular to the linkers. In addition, the outer linkers are more accessible on a 2D surface, allowing invader strands diffusing on the mica to engage them more readily. Together, these factors could make the outer linkers more susceptible to initial displacement, facilitating their detachment once an invader engages the toehold. This observation also highlights the importance of the placement of the toehold-containing helix within the nanostructure. In 3D origami, if the toehold helix was embedded within the core of the structure, it can be harder to access, which may ultimately reduce the efficiency of the TMSD-driven nanomechanical response.

Our HS-AFM data shows that the complete separation of the dimer proceeds through a series of transient linker disconnections (Figure 3). When all 5 linkers are intact, the dimer forms a multivalently connected state, in which the left and right monomers are held in close proximity. Disconnection of a single linker does not substantially alter this geometry, as the remaining linkers maintain a high local effective concentration of the complementary toehold linker between the monomers. As a result, a broken connection is likely to reform rapidly, giving rise to short-lived and reversible disconnection events. Furthermore, the exponential distribution of the transient disconnection states (Figure 3e) indicates its stochastic nature, the rate of re-connection is independent from the duration of the disconnected state. These transient disconnections are initiated by stochastic encounters between invader strands and the linkers’ toeholds and are governed by invader diffusion. Importantly, the probability of a disconnection event does not depend on the cumulative number of prior disconnection and re-connection events experienced by a given linker. Our approach thus reveals that TMSD in this multivalent system is dominated by stochastic, kinetically driven fluctuations between connected and disconnected linker states within a confined geometry, rather than by a smooth, effectively irreversible stepwise progression towards separation, as is typically observed for freely diffusing strands in solution, where re-hybridisation after displacement is entropically not favoured^19,64,65,72^.

Overall, the TMSD-driven separation proceeds through repeated, transient disconnection events until successive loss of multiple linkers, once sufficient linkers are lost in close succession, the remaining connections can no longer maintain a high local effective concentration of the complementary toehold linker between the monomers, leading to a rapid, cooperative separation of the dimer. After the full, permanent separation of all 5 linkers, the resulting monomers could not reconnect and form the dimer again. Free monomers could randomly and weakly interact with another monomer for a very small fraction of time, these interactions were unstable and could be easily identified through the orientation marker.

### A multiplexable miRNA sensor

Since both DNA and RNA invaders efficiently triggered TMSD based separation of the dimer, the linker sequences could be replaced with target miRNA sequences, thus utilising the dimer as miRNA sensor. Accordingly, we modified the dimer’s linker sequences to create a miRNA-responsive dimer capable of detecting miRNA in solution. (Figure 4a). As a proof of concept, we selected miR-532-5p, a miRNA associated with traumatic brain injury (TBI)^32^ , as our target. The DNA sequence of the linker was engineered so that miR-532 could act as an invader, triggering the separation of the dimer via strand displacement. The successful construction of the miR-532-responsive dimer and its displacement mechanism were validated using gel electrophoresis and AFM (Figure S15a).

**Figure 4.**
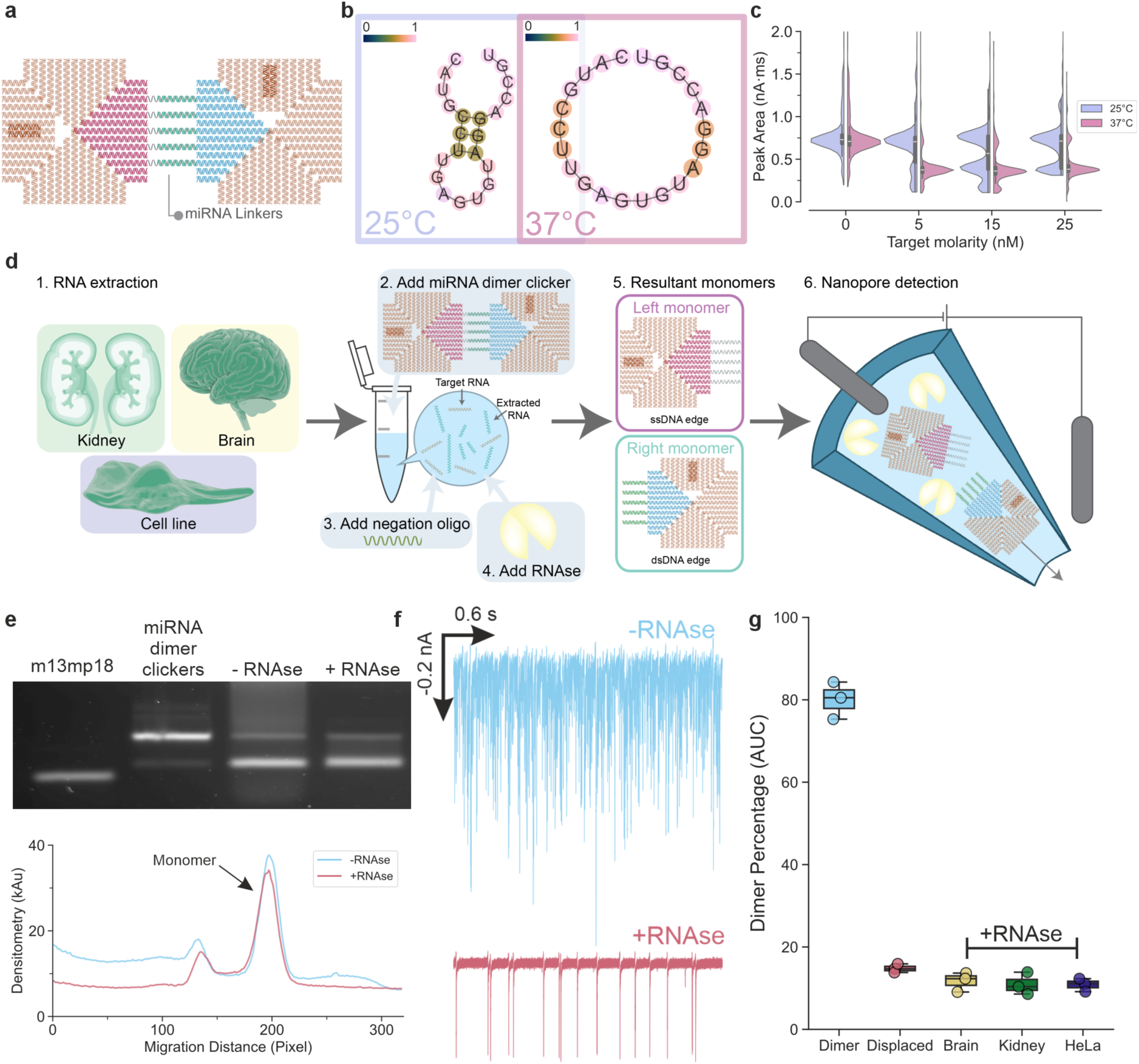
Dimer for the detection of miR-532. (**a**) Schematic illustration of the dimer specific for miR-532. The miR-532-5p (5’ arm of the miRNA) sequence was used to construct the miRNA linker dimer. (**b**) The minimum free energy (MFE) prediction of the miR-532-5p secondary structure at 10 mM Na^+^ condition under either 25°C or 37°C with ViennaRNA 2.0 package. The colour gradient showed the probability of either pairing (if paired) or unpairing (if unpaired) with 0 being the lowest probability and 1 being the highest probability. The miR-532 adopted an open conformation (no secondary structure) at 37°C contrasting its hairpin secondary structure at 25°C. (**c**) The peak area of the nanopore translocation events of the miR-532 dimer at 1 nM. The data showed that the displacement carried out at 25°C leads to inefficient separation of the miR-532 dimer. The displacement was more efficient when carried out at 37°C where 5 nM was sufficient to separate the dimer into monomers (14% dimer remaining), in contrast to 5 nM at 25°C where 80% of the dimer remained as dimer. (**d**) Schematic illustration of the selective detection under RNAse contaminated environment. Tissue or cell line extracted RNAs are used as RNA background, the RNA background was spiked with miR-532 (target RNA) at final concentration of 5 nM. The miR-532 responsive dimer (1 nM) was added to the mixture and followed by the addition of negation ssDNA oligo, and then RNAse. The resultant RNAse treated samples were quantified with nanopore. (**e**) The agarose gel analysis on the dimer treated with or without RNAse after separation. Larger smearing background could be observed in the densitometry when not treated with RNAse. (**f**) The nanopore translocation trace of the samples with or without the RNAse treatment, due to the high quantity of the background RNAs translocation through the nanopore, the displaced origami monomer signals were hidden. The RNAse treated samples revealed a clean trace showing the translocation of the displaced origami monomers. (**g**) The detection of the displaced dimer at normal pure buffer condition and RNA background with RNAse contaminated environment.

Due to sequence differences between miR-532 and the original artificial invader, we observed that miR-532 was less efficient at displacing the dimer. While the artificial RNA sequence achieved complete dimer displacement at 15 nM and 25°C (Figure 2), miR-532 only displaced about 50% of the dimer under the same conditions (Figure 4c). We hypothesize that the secondary structure of miR-532 may hinder the TMSD reaction in this context. Using the ViennaRNA package^80^, we calculated the minimum free energy (MFE) and the secondary structure under lower salt condition (Figure 4b). The simulation indicated that at 25°C, miR-532 adopted a hairpin secondary structure with only 4 nt accessible to the linker’s toehold. This condition was less thermodynamically favourable for the TMSD reaction to proceed efficiently. We also analysed the secondary structure of the artificial sequence (Figure S15b) and found no stable or defined secondary structure, suggesting it could fully hybridize with the linker’s toehold, thereby promote efficient TMSD. Indeed, previous studies have shown that RNA invaders with a 5’ toehold binding region tend to form RNA:RNA duplexes at the 3’ end of the target, which significantly slows the TMSD reaction down and reduces its overall efficiency^71^.

By increasing the incubation temperature from 25°C to 37°C, the secondary hairpin structures of miR-532 were disrupted (Figure 4b, Supporting Table 1). We confirmed that the miR-532-responsive dimers remained stable at 37°C for at least 30 minutes (Figure S15c,d). TMSD reactions were then performed at 37°C for 30 minutes, resulting in near complete separation (14% dimer remaining) of 1 nM miR-532 dimer with as little as 5 nM miR-532, as determined by nanopore analysis (Figure 4c). These results contrast sharply with the results at 25°C, where 80% of dimers remained intact at 5 nM miR-532 (Figure 4c). These findings suggest that disrupting the secondary structure of the invader strand through temperature modulation significantly enhances the reactivity, and the efficiency of the dimer displacement reaction.

Previous studies using nanopore-based miRNA detection from fluid have typically relied on the miRNA itself serves as a linker between two larger nano-entities like nanoparticles or multi-armed DNA nanostars^45,57^. This concept can be replicated using our monomer origamis, with the miR-532 oligo acting as a bridge to hybridize and connect two tiles to form a dimer (Figure S16). However, this approach has several disadvantages: it is entropically unfavourable to form a higher order structure; the miR-532 linked dimer must be kept in RNAse free or inactivated environment, because of the fast degradation of RNA when exposed to RNAse^36^; it could lead to the formation of unintended higher order structures that require removal before nanopore readout. As demonstrated, the miR-532-linked dimer is highly sensitive to RNase exposure and can be readily dissociated into monomers (Figure S16b). This introduces a significant source of uncertainty in assays aiming to quantify miRNA concentrations, as RNase contamination could lead to false negatives. Consequently, this method requires specialized training and stringent storage conditions, including -80°C storage for RNA, to ensure reliable detection.

In contrast, our dimer approach is designed to be irreversible after the TMSD reaction and can be stored at 4°C (and potentially room temperature) by simply incorporating the addition of DNA negation strands to hybridise with the right monomers after TMSD reaction. This step prevents the re-hybridisation between the left and right monomers after the RNAse digested the RNA:ssDNA duplex (Figure 4d). We introduced a counter intuitive step to add RNAse prior to the analysis, to remove any background RNA (including the target miRNAs) that would interfere with the nanopore readout^41,47^, and to simulate the quantification of miRNA assay in an RNAse contaminated environment. At this stage, the TMSD reactions had already proceeded irreversibly, meaning that the removal of RNA, including the target miRNAs, no longer affecting the overall dimer population.

To demonstrate the effectiveness of our approach, we supplemented total RNA extracted from tissues or cell lines with 5 nM of synthetic miR-532 (Figure 4d). A 1 nM concentration of the miR-532-responsive dimer was then added to the RNA mixture and incubated at 37°C for 30 minutes. This was followed by the addition of negation oligos (short DNA strands complementary to the ssDNA linker) of the right monomer, then a second 30-minute incubation at 37°C. Finally, the samples were treated with RNase (50 U/mL RNase A and 2,000 U/mL RNase T1) at 25°C for 30 minutes. Gel electrophoresis and densitometry confirmed that the miR-532 dimers were successfully separated into monomers, which migrated to the expected positions on the gel (Figure 4e, Figure S17). In contrast, samples that did not undergo RNase treatment showed significant background smearing, indicating the presence of high RNA content (∼100 ng). This background RNA interfered with nanopore analysis, producing numerous translocation peaks from both DNA origami structures (monomers and dimers) and the background RNA. As a result, the distinct peak signatures of the DNA origami were obscured (Figure 3f). However, RNase digestion effectively removed this interference, allowing clear detection of the displaced monomers via nanopore analysis (Figure 3g). This confirmed the presence of at least 5 nM of miR-532-5p in all RNA extract samples.

Our DNA origami design enables high spatial addressability where each linker can target a specific miRNA sequence (Figure 5a). We selected 4 additional miRNAs-5p, the cancer and cardiovascular diseases associated miR-21^81^, viral infection associated let-7a^82^, and the TBI associated miR-221 and miR-629^83^. The formation of this multiplex dimer follows the same procedure as the formation of the miRNA dimer and the original artificial sequence dimer, and the resultant dimer could be seen in Figure 5b. To initiate the dimer disassembly, a mixture of five miRNA invaders (miR-221, miR-629, miR-21, let-7a, and miR-532) was applied at a final concentration of 5 nM and incubated at 37 °C. The dimers were incubated with different mixtures of invaders form by systematically removing an individual miRNA from the mixture to test the minimum number of invaders needed to achieve the disassembly (Figure 5c). The percentage of dimers reduces as the number of invaders increased in the mix until 4 invaders are added (Figure 5d).

**Figure 5.**
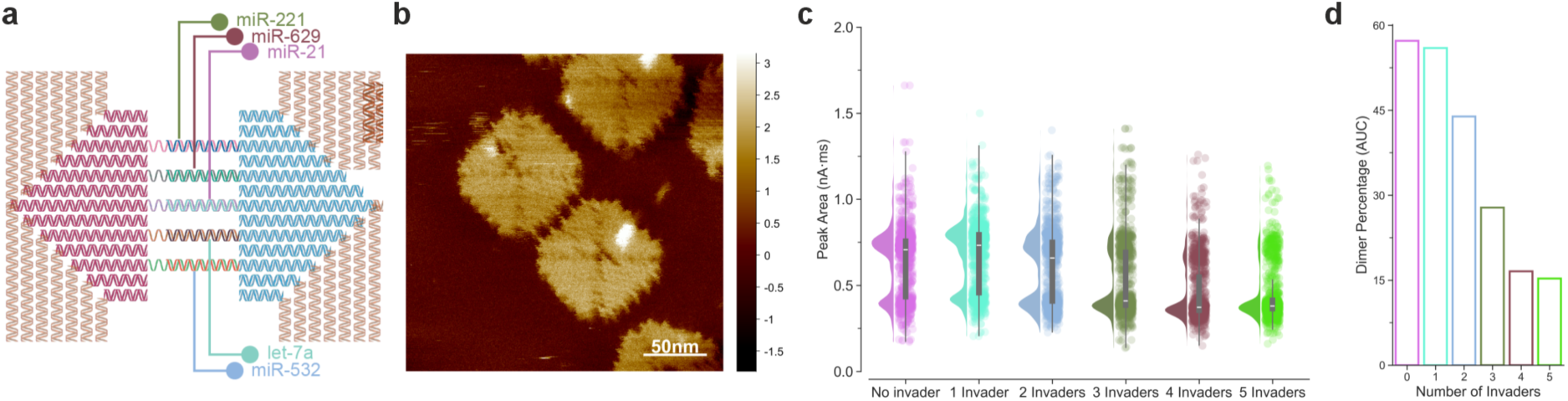
Multiplex miRNA detection. (**a**) The formation of the multiplex miRNA dimer with different sequences of miRNA (miR-221, miR-629, miR-21, let-7a and miR-532) as linkers. (**b**) AFM images of the formation of the multiplex miRNA dimer. (**c**) The displacement of the different linkers’ dimer clickers quantified with nanopore (n=1000). The number of invaders correspond to the species of the miRNA, 5 invaders mean a cocktail combination of miR-221, miR-629, miR-21, let-7a and miR-532 while 1 invader means only miR-532 was added at 5 nM. (**d**) Quantification of the dimer percentage.

These proof-of-concept results demonstrate that our approach can be employed for the multiplexed detection of three miRNAs. Full dimer separation (dimer percentage less than 15%) occurs when only two linkers are left, irrespective of whether the invader strand is added or not, a finding further confirmed by HS-AFM (Figure 3). The progressive decrease in dimer percentage with one to four invaders reflects the gradual reduction in stability of the connections between the monomers.

The simultaneous presence of multiple miRNAs has been explored for early-stage disease detection, including cancers, with several studies highlighting the value of disease detection using miRNA panels comprising at least five or more upregulated miRNAs in body fluids^84–86^. Such multiplexed signatures have also been shown to discriminate malignant from benign tissues and to provide additional information for classifying disease subtypes^86^. Our design ensures that the dimer disassembles only when all the target miRNAs are upregulated, additionally, we have already demonstrated that up to nine linkers can be incorporated into our DNA origami design (Figure S1a), indicating that, at least seven miRNAs could be monitored within a single nanopore based assay.

## Conclusion

The use of polymer electrolyte as the trans chamber electrolyte allowed us to perform detection at materials native buffer condition, it was widely accepted that for translocation experiment with solid-state nanopore, a minimum of 0.1 M salt (reported up 4 M)^87,88^ would be needed for the system to be sensitive to the translocation of materials. Here, against expectations, the detections were carried out with maximum 15 mM of divalent ions (Mg^2+^). The avoidance of additional high concentration of salt reduced the physical modification on the DNA itself, as concentration of salt modified the properties of the DNA^89,90^, which could potentially affect the translocation signals generated^68^. Further development in this could potentially allow solid-state nanopore to be a tool to probe DNA physical properties.

We utilise the established TMSD mechanic to disassemble the dimerised DNA origami with either DNA or RNA and subsequently showed that exposure of RNAse to the DNA origami did not interfere with the detection of the already separated DNA origami, the population differences were readily quantified with our nanopore system as a potential point of care biosensors, bypassing laborious procedures in sample handling in potentially contaminated environment. This approach here only required the initial RNA and DNA origami preparation to be clean, significantly reduced the risk of RNA post extraction degradation due to mishandling^91^. Our approach utilised RNAse to digest irrelevant background RNA after the TMSD step act to reveal the true DNA origami translocation signal and enabled multi-miRNA detections through a multiplex approach. This strategy could be further refined and implemented into the nanopore sequencing device like MinION, providing an alternative strategy for miRNA detection alongside the recently established method^39^.

Our detection strategy relies on a rather simple separation of masses, unlike a recent approach that relies on structural conformational change^58^, subsequently a faster and more responsive assay. The 2D nature of the origami designs allowed us to trace and visualise the molecular motion of the TMSD in action with HS-AFM in real time. Upon the introduction of the invader strands, the five DNA linkers connecting the dimer separated at different time points, with some linkers temporarily reconnecting before complete dissociation. Kymograph analysis of AFM height profiles enabled state tracking (connected/disconnected) of each linker, revealing detailed dynamic behaviour of the linkers and general separation of the dimer requires less than 20 seconds once initiated by the invaders. The outer linkers (L1 and L5) were the first to separate in most cases, likely due to their greater exposure on the mica surface. Interestingly, HS-AFM revealed that dimer separation proceeds through multiple short-lived and reversible linker disconnection–reconnection events, followed by a rapid and irreversible separation once the remaining linkers can no longer maintain close spatial confinement between the monomers. These observations show that linker position and nanostructure geometry govern the TMSD response, and establish HS-AFM as a powerful tool for probing dynamic DNA nanomechanics in real time.

## Methods

For full detailed method, please refer to the supporting information

### DNA origami folding and dimerization

All DNA strands used throughout this study were purchased from Integrated DNA Technologies with standard desalting. The caDNAno (2.4.7) software was used to design the DNA origami structures^5^. All origamis were folded using the 7249 nt M13mp18 circular single-stranded DNA scaffold (type p7249; tilibit). All folded origami were stored at 4°C or immediately purified. The caDNAno files of the origami designs list of staples could be found in the repository associated with this publication.

For all purposes unless otherwise specified, the origami buffer is composed of 10 mM Tris-Acetate, 15 mM Mg-Acetate, 1 mM EDTA, pH 8.0.

For a single folding reaction of 80 µl, the folding mixture contained: 10 nM M13mp18 scaffold, 12.5 mM magnesium acetate, 75 nM bridge mixture, 75 nM interior mixture, 75 nM edge mixture, and diluted in 10 mM tris-HCl, 1mM Ethylenediaminetetraacetic acid (EDTA) at pH 8.0 (1× TE buffer). The folding of the DNA origami tiles was carried out within a thermal cycler by heating the folding mixture to 90°C for 2 minutes, followed by a gradual temperature decrease from 90°C to 20°C at the rate of 1°C per minute. The temperature was then further decreased to 4°C until use.

The purification of the folded monomers followed our published method using solid-phase reversible immobilization (SPRI) beads^92^ with a modification. The HighPrep™ PCR Clean-up System (AC-60050; Magbio) SPRI beads was used, the buffer suspending the SPRI beads were exchanged to 2.5% (w/v) Poly(ethylene) glycol (PEG) 8K, 500 mM MgCl_2_ via pelleting the SPRI beads down and resuspending the beads in the custom buffer, the concentration of the beads to buffer was at 20 mg/ml. The buffer modification improved the performance of the SPRI beads for the purpose of DNA origami purification.

After the folding reaction, the reaction was diluted to 100 µl with DNA origami buffer, then 40 µl (0.4× beads to solution volume ratio) of the buffer modified SPRI beads were added to the solution and mix via pipetting until homogenous colour and incubated for 5 minutes. Then, the SPRI beads were separated with a 3D-printed magnetic rack, the solution was aspirated and discarded. The pellet was washed with 200 µl of 80% (v/v) ethanol once, after the wash, the pellet was resuspended in 40 µl of the DNA origami buffer, incubated for 5 minutes (40 µl was used to concentrate the DNA origami). The SPRI beads were separated from the solution once again with the magnetic rack, the purified DNA origami solution was aspirated and collected to a PCR tube, the tube was ready for thermal de-clump step. The PCR tube was heated to 50°C using a thermal cycler for 2 mins, followed by a gradual decrease of the temperature to 20°C at the rate of 3°C/min. The temperature was then further decreased down to 4°C and stored at 4°C until use.

The yield of the DNA origami after the SPRI purification was checked, to generate the asymmetric dimer , the left and right monomer was mixed in the same PCR tube so that the final mixture contains 400 ng of left monomers and 400 ng of right monomers in a total volume of 40 µl per PCR tube (multiple tubes of the same mixture would be used if large volume of dimer is needed). The mixture was then heated to 55°C and the temperature was then gradually decreased to 20°C at a 1°C/min rate. The temperature was then further decreased down to 4°C until use.

### Atomic Force Microscopy (AFM)

Freshly cleaved mica discs were pre-treated with 5 µl of 10 mM NiCl_2_, then immediately 10 µl of the DNA origami samples were deposited onto the mica by pipetting directly onto the NiCl_2_ droplet, then the samples were topped up with 100 µl of DNA origami buffer and incubated for 5 minutes. The mica was then washed twice with the origami buffer, topped with 100 µl of origami buffer for liquid imaging. The DNA origami samples were imaged using a Bruker Dimension Fastscan Bio (Santa Barbara, CA, USA) with PEAKFORCE-HIRS-F-B probes with a small laser spot on the Fastscan head. The imaging was carried out via PeakForce tapping with ScanAsyst™ liquid imaging mode via the Nanoscope software. All images acquired have a pixel resolution of minimum 512 × 512 to 1024 × 1024, and the images were analysed with NanoLocz AFM image analysis software (v1.20)^63^.

### High-Speed Atomic Force Microscopy (HS-AFM)

All high speed-atomic force microscopy (HS-AFM) measurements were performed using a NanoRacer HS-AFM (Bruker, Germany) instrument in amplitude modulation mode. All HS-AFM measurements were obtained in liquid and ambient temperature in an acoustic isolation housing on an active antivibration table using short cantilevers (USC-F1.2-k0.15, NanoWorld, Switzerland) with nominal spring constants of 0.15 N·m^−1^, resonance frequencies of ∼0.6 MHz and quality factors of ∼2. To prepare the sample for HS-AFM imagining of DNA Origami, freshly cleaved mica was treated with 5 µl of 10 mM Ni^2+^/Mg^2+^ salt solution, to promote the adsorption of DNA origami to the mica. Thereafter, 2-4 μl of DNA Origami sample was incubated on the mica and left for incubation for 2-3 minutes before rinsing, with 5 via fluid exchange with 5 µl of buffer solution. The sample holder was then filled with 1 ml with imaging buffer (5 mM Tris-HCl (pH 8.0), 1 mM EDTA, 20 mM MgCl_2_ and 5 mM NaCl). To visualize TMSD, 10 µl of the invader strands at a concentration of 1 µM were added to the imaging buffer. The images and videos were analysed with NanoLocz image analysis software (v1.30)^63^.

### Nanopore measurement

Quartz capillaries of 1.0 mm outer diameter and 0.5 mm inner diameter (QF100-50-7.5; Sutter Instrument) were used to fabricate the glass nanopores using the SU-P2000 laser puller (World Precision Instruments). A two-line protocol was used: line 1, HEAT 750/FIL 4/VEL 30/DEL 150/PUL 80, followed by line 2, HEAT 700/FIL 3/VEL 40/DEL 135/PUL 180. The pulling protocol was instrument specific, and these would vary between different laser pullers.

The generation of the polymer electrolyte bath followed previously published method ^41^. To generate 40 ml of the polymer electrolyte with a composition of 50% (w/v) PEG 35K, 0.1 M KCl, 4 ml of 1 M KCl was mixed with 16 ml of ddH_2_O and 20 g of PEG 35K (81310; Sigma) inside a 50 ml centrifuge tube. The tube was then incubated at 85°C for 2 hours and left over night at 37°C, electrolyte was then aliquoted inside multiple 7 ml Bijou tubes, stored on shelves away from sunlight at room temperature (E1412-0710; Starlab), the aliquot would be discarded after 1 month since the first immersion of the nanopore and 6 months after the initial generation.

The translocation experiment followed a similar procedure from the previous publications^16,40–42,92,93^ The glass nanopores were all filled with 9.4 ng/µl of origami (4 nM for monomers, 2 nM for dimer), all origami were diluted in the origami buffer without further addition of any salt. In each measurement, the glass nanopore was fitted with an Ag/AgCl working electrode and immersed into the polymer electrolyte bath with an Ag/AgCl reference electrode. The ionic current trace was recorded using a MultiClamp 700B patch-clamp amplifier (Molecular Devices) in voltage-clamp mode. The signal was filtered using a low-pass filter at 20 kHz unless specified, digitized with a Digidata 1550B at a 100 kHz (10 μs) sampling rate, and recorded using the software pClamp 10 (Molecular Devices). Unless otherwise specified, all translocations were carried out at -500 mV.

The python code used to perform the translocation event detection could be accessed at https://github.com/chalmers4c/Nanopore_event_detection. For all events calling, a threshold line of 7 sigmas away from baseline is used.

## Data availability

The AFM images and HS-AFM video, translocation trace data, DNA origami design, staples list and relevant sequences from this work can be freely accessed via the University of Leeds data repository (https://doi.org/10.5518/1770).

## Supporting Information

Supporting information contains additional detailed methods and Supporting Figures.

## Author Contributions

C.C.C.C. and V.G. designed and performed the experiments. C.C.C.C. illustrated all schematics. G.R.H., C.W. and P.A. helped with data analysis and acquired the funding. All authors wrote and corrected the manuscript.

## Acknowledgement

We thank the group of bioelectronics member for providing insightful feedback. We thank Dr Alexander Kulak of University of Leeds for imaging the nanopore with the scanning electron microscopy. C.C.C.C. thanks for an inspiring talk with Dr Fabio Marcuccio from Human Technopole.

## References

1 Seeman, N., Sleiman, H. DNA nanotechnology. Nat Rev Mater 3, doi:10.1038/natrevmats.2017.68 (2018).

2 Winfree, E., Liu, F., Wenzler, L. A. & Seeman, N. C. Design and self-assembly of two-dimensional DNA crystals. Nature 394, 539–544, doi:10.1038/28998 (1998).

3 Rothemund, P. W. K. Folding DNA to create nanoscale shapes and patterns. Nature 440, 297–302, doi:10.1038/nature04586 (2006).

4 Dey, S. et al. DNA origami. Nature Reviews Methods Primers 1, doi:10.1038/s43586-020-00009-8 (2021).

5 Douglas, S. M. et al. Rapid prototyping of 3D DNA-origami shapes with caDNAno. Nucleic Acids Research 37, 5001–5006, doi:10.1093/nar/gkp436 (2009).

6 Posnjak, G. et al. Diamond-lattice photonic crystals assembled from DNA origami. Science 384, 781–785, doi:10.1126/science.adl2733 (2024).

7 Liu, H. et al. Inverse design of a pyrochlore lattice of DNA origami through model-driven experiments. Science 384, 776–781, doi:10.1126/science.adl5549 (2024).

8 Woo, S. & Rothemund, P. W. K. Programmable molecular recognition based on the geometry of DNA nanostructures. Nature Chemistry 3, 620–627, doi:10.1038/nchem.1070 (2011).

9 He, Z., Shi, K., Li, J. & Chao, J. Self-assembly of DNA origami for nanofabrication, biosensing, drug delivery, and computational storage. iScience 26, 106638, doi:10.1016/j.isci.2023.106638 (2023).

10 Wang, J., Zhou, Z., Yue, L., Wang, S. & Willner, I. Switchable Triggered Interconversion and Reconfiguration of DNA Origami Dimers and Their Use for Programmed Catalysis. Nano Letters 18, 2718–2724, doi:10.1021/acs.nanolett.8b00793 (2018).

11 Wu, N. & Willner, I. Programmed dissociation of dimer and trimer origami structures by aptamer–ligand complexes. Nanoscale 9, 1416–1422, doi:10.1039/c6nr08209b (2017).

12 Wu, N. & Willner, I. DNAzyme-Controlled Cleavage of Dimer and Trimer Origami Tiles. Nano Letters 16, 2867–2872, doi:10.1021/acs.nanolett.6b00789 (2016).

13 Wu, N. & Willner, I. pH-Stimulated Reconfiguration and Structural Isomerization of Origami Dimer and Trimer Systems. Nano Letters 16, 6650–6655, doi:10.1021/acs.nanolett.6b03418 (2016).

14 Zhou, Z. et al. Triggered Dimerization and Trimerization of DNA Tetrahedra for Multiplexed miRNA Detection and Imaging of Cancer Cells. Small 17, doi:10.1002/smll.202007355 (2021).

15 Padilla, J. E., Liu, W. & Seeman, N. C. Hierarchical self assembly of patterns from the Robinson tilings: DNA tile design in an enhanced Tile Assembly Model. Natural Computing 11, 323–338, doi:10.1007/s11047-011-9268-7 (2011).

16 Confederat, S., Sandei, I., Mohanan, G., Wälti, C. & Actis, P. Nanopore fingerprinting of supramolecular DNA nanostructures. Biophysical Journal 121, 4882–4891, doi:10.1016/j.bpj.2022.08.020 (2022).

17 Tikhomirov, G., Petersen, P. & Qian, L. Fractal assembly of micrometre-scale DNA origami arrays with arbitrary patterns. Nature 552, 67–71, doi:10.1038/nature24655 (2017).

18 Tikhomirov, G., Petersen, P. & Qian, L. Programmable disorder in random DNA tilings. Nature Nanotechnology 12, 251–259, doi:10.1038/nnano.2016.256 (2016).

19 Srinivas, N. et al. On the biophysics and kinetics of toehold-mediated DNA strand displacement. Nucleic Acids Research 41, 10641–10658, doi:10.1093/nar/gkt801 (2013).

20 Yurke, B., Turberfield, A. J., Mills, A. P., Simmel, F. C. & Neumann, J. L. A DNA-fuelled molecular machine made of DNA. Nature 406, 605–608, doi:10.1038/35020524 (2000).

21 Simmel, F. C., Yurke, B. & Singh, H. R. Principles and Applications of Nucleic Acid Strand Displacement Reactions. Chemical Reviews 119, 6326–6369, doi:10.1021/acs.chemrev.8b00580 (2019).

22 Chandrasekaran, A. R. et al. Cellular microRNA detection with miRacles: microRNA-activated conditional looping of engineered switches. Science Advances 5, doi:10.1126/sciadv.aau9443 (2019).

23 Kim, M. et al. Harnessing a paper-folding mechanism for reconfigurable DNA origami. Nature 619, 78–86, doi:10.1038/s41586-023-06181-7 (2023).

24 Kocabey, S., Chiarelli, G., Acuna, G. P. & Ruegg, C. Ultrasensitive and multiplexed miRNA detection system with DNA-PAINT. Biosensors and Bioelectronics 224, 115053, doi:10.1016/j.bios.2022.115053 (2023).

25 Chandrasekaran, A. R. et al. DNA nanotechnology approaches for microRNA detection and diagnosis. Nucleic Acids Research 47, 10489–10505, doi:10.1093/nar/gkz580 (2019).

26 Wang, D. et al. Molecular Logic Gates on DNA Origami Nanostructures for MicroRNA Diagnostics. Analytical Chemistry 86, 1932–1936, doi:10.1021/ac403661z (2014).

27 Kuzuya, A., Sakai, Y., Yamazaki, T., Xu, Y. & Komiyama, M. Nanomechanical DNA origami ’single-molecule beacons’ directly imaged by atomic force microscopy. Nature Communications 2, doi:10.1038/ncomms1452 (2011).

28 Winter, J., Jung, S., Keller, S., Gregory, R. I. & Diederichs, S. Many roads to maturity: microRNA biogenesis pathways and their regulation. Nature Cell Biology 11, 228–234, doi:10.1038/ncb0309-228 (2009).

29 O’Brien, J., Hayder, H., Zayed, Y. & Peng, C. Overview of MicroRNA Biogenesis, Mechanisms of Actions, and Circulation. Frontiers in Endocrinology 9, doi:10.3389/fendo.2018.00402 (2018).

30 Bushati, N. & Cohen, S. M. microRNA Functions. Annual Review of Cell and Developmental Biology 23, 175–205, doi:10.1146/annurev.cellbio.23.090506.123406 (2007).

31 Skommer, J. et al. Small molecules, big effects: the role of microRNAs in regulation of cardiomyocyte death. Cell Death & Disease 5, e1325–e1325, doi:10.1038/cddis.2014.287 (2014).

32 Atif, H. & Hicks, S. D. A Review of MicroRNA Biomarkers in Traumatic Brain Injury. Journal of Experimental Neuroscience 13, 117906951983228, doi:10.1177/1179069519832286 (2019).

33 Jeter, C. B. et al. Biomarkers for the Diagnosis and Prognosis of Mild Traumatic Brain Injury/Concussion. Journal of Neurotrauma 30, 657–670, doi:10.1089/neu.2012.2439 (2013).

34 Mavroudis, I. et al. The Role of Microglial Exosomes and miR-124-3p in Neuroinflammation and Neuronal Repair after Traumatic Brain Injury. Life 13, 1924, doi:10.3390/life13091924 (2023).

35 Xu, D. et al. MicroRNAs in extracellular vesicles: Sorting mechanisms, diagnostic value, isolation, and detection technology. Frontiers in Bioengineering and Biotechnology 10, doi:10.3389/fbioe.2022.948959 (2022).

36 Green, M. R. & Sambrook, J. How to Win the Battle with RNase. Cold Spring Harbor Protocols 2019, pdb.top101857, doi:10.1101/pdb.top101857 (2019).

37 Chandrasekaran, A. R., Trivedi, R. & Halvorsen, K. Ribonuclease-Responsive DNA Nanoswitches. Cell Reports Physical Science 1, 100117, doi:10.1016/j.xcrp.2020.100117 (2020).

38 Siddika, T. & Heinemann, I. U. Bringing MicroRNAs to Light: Methods for MicroRNA Quantification and Visualization in Live Cells. Front Bioeng Biotechnol 8, 619583, doi:10.3389/fbioe.2020.619583 (2020).

39 Koch, C. et al. Nanopore sequencing of DNA-barcoded probes for highly multiplexed detection of microRNA, proteins and small biomarkers. Nature Nanotechnology 18, 1483–1491, doi:10.1038/s41565-023-01479-z (2023).

40 Chau, C. C., Radford, S. E., Hewitt, E. W. & Actis, P. Macromolecular Crowding Enhances the Detection of DNA and Proteins by a Solid-State Nanopore. Nano Letters 20, 5553–5561, doi:10.1021/acs.nanolett.0c02246 (2020).

41 Chau, C. et al. Probing RNA Conformations Using a Polymer–Electrolyte Solid-State Nanopore. ACS Nano 16, 20075–20085, doi:10.1021/acsnano.2c08312 (2022).

42 Marcuccio, F. et al. Mechanistic Study of the Conductance and Enhanced Single-Molecule Detection in a Polymer–Electrolyte Nanopore. ACS Nanoscience Au 3, 172–181, doi:10.1021/acsnanoscienceau.2c00050 (2023).

43 Confederat, S. et al. Next-Generation Nanopore Sensors Based on Conductive Pulse Sensing for Enhanced Detection of Nanoparticles. Small 20, doi:10.1002/smll.202305186 (2023).

44 Xue, L. et al. Solid-state nanopore sensors. Nature Reviews Materials 5, 931–951, doi:10.1038/s41578-020-0229-6 (2020).

45 Ren, R. et al. Single-Molecule Binding Assay Using Nanopores and Dimeric NP Conjugates. Advanced Materials 33, doi:10.1002/adma.202103067 (2021).

46 Cai, S. et al. Single-molecule amplification-free multiplexed detection of circulating microRNA cancer biomarkers from serum. Nature Communications 12, doi:10.1038/s41467-021-23497-y (2021).

47 Bošković, F. & Keyser, U. F. Nanopore microscope identifies RNA isoforms with structural colours. Nature Chemistry 14, 1258–1264, doi:10.1038/s41557-022-01037-5 (2022).

48 Bell, N. A. W. & Keyser, U. F. Digitally encoded DNA nanostructures for multiplexed, single-molecule protein sensing with nanopores. Nature Nanotechnology 11, 645–651, doi:10.1038/nnano.2016.50 (2016).

49 Dorey, A. & Howorka, S. Nanopore DNA sequencing technologies and their applications towards single-molecule proteomics. Nat Chem 16, 314–334, doi:10.1038/s41557-023-01322-x (2024).

50 Chau, C. C. C., Weckman, N. E., Thomson, E. E. & Actis, P. Solid-State Nanopore Real-Time Assay for Monitoring Cas9 Endonuclease Reactivity. ACS Nano 19, 3839–3851, doi:10.1021/acsnano.4c15173 (2025).

51 Sandler, S. E. et al. Sensing the DNA-mismatch tolerance of catalytically inactive Cas9 via barcoded DNA nanostructures in solid-state nanopores. Nature Biomedical Engineering 8, 325–334, doi:10.1038/s41551-023-01078-2 (2023).

52 Charron, M., Briggs, K., King, S., Waugh, M. & Tabard-Cossa, V. Precise DNA Concentration Measurements with Nanopores by Controlled Counting. Analytical Chemistry 91, 12228–12237, doi:10.1021/acs.analchem.9b01900 (2019).

53 Raveendran, M., Lee, A. J., Wälti, C. & Actis, P. Analysis of 2D DNA Origami with Nanopipettes. ChemElectroChem 5, 3014–3020, doi:10.1002/celc.201800732 (2018).

54 Raveendran, M., Lee, A. J., Sharma, R., Wälti, C. & Actis, P. Rational design of DNA nanostructures for single molecule biosensing. Nature Communications 11, doi:10.1038/s41467-020-18132-1 (2020).

55 Yan, H. et al. Central Limit Theorem-Based Analysis Method for MicroRNA Detection with Solid-State Nanopores. ACS Applied Bio Materials 4, 6394–6403, doi:10.1021/acsabm.1c00587 (2021).

56 Zahid, O. K., Wang, F., Ruzicka, J. A., Taylor, E. W. & Hall, A. R. Sequence-Specific Recognition of MicroRNAs and Other Short Nucleic Acids with Solid-State Nanopores. Nano Letters 16, 2033–2039, doi:10.1021/acs.nanolett.6b00001 (2016).

57 Beamish, E., Tabard-Cossa, V. & Godin, M. Digital counting of nucleic acid targets using solid-state nanopores. Nanoscale 12, 17833–17840, doi:10.1039/d0nr03878d (2020).

58 Long, L. et al. Reconfigurable DNA origami hinges for nanopore detection of microRNA. Nano Research, doi:10.26599/nr.2025.94907604 (2025).

59 Chau, C., Mohanan, G., Macaulay, I., Actis, P. & Wälti, C. Automated Purification of DNA Origami with SPRI Beads. Small 20, doi:10.1002/smll.202308776 (2023).

60 Ando, T. et al. A high-speed atomic force microscope for studying biological macromolecules. Proceedings of the National Academy of Sciences 98, 12468–12472, doi:10.1073/pnas.211400898 (2001).

61 Gilmore, J. L. et al. Single-Molecule Dynamics of the DNA−EcoRII Protein Complexes Revealed with High-Speed Atomic Force Microscopy. Biochemistry 48, 10492–10498, doi:10.1021/bi9010368 (2009).

62 Heath, G. R. et al. Localization atomic force microscopy. Nature 594, 385–390, doi:10.1038/s41586-021-03551-x (2021).

63 Heath, G. R., Micklethwaite, E. & Storer, T. M. NanoLocz: Image Analysis Platform for AFM, High-Speed AFM, and Localization AFM. Small Methods, doi:10.1002/smtd.202301766 (2024).

64 Zhang, D. Y. & Winfree, E. Control of DNA Strand Displacement Kinetics Using Toehold Exchange. Journal of the American Chemical Society 131, 17303–17314, doi:10.1021/ja906987s (2009).

65 Liu, H. et al. Kinetics of RNA and RNA:DNA Hybrid Strand Displacement. ACS Synthetic Biology 10, 3066–3073, doi:10.1021/acssynbio.1c00336 (2021).

66 Wang, B., Thachuk, C., Ellington, A. D., Winfree, E. & Soloveichik, D. Effective design principles for leakless strand displacement systems. Proceedings of the National Academy of Sciences 115, doi:10.1073/pnas.1806859115 (2018).

67 SantaLucia, J. & Hicks, D. The Thermodynamics of DNA Structural Motifs. Annual Review of Biophysics and Biomolecular Structure 33, 415–440, doi:10.1146/annurev.biophys.32.110601.141800 (2004).

68 He, L. et al. DNA origami characterized via a solid-state nanopore: insights into nanostructure dimensions, rigidity and yield. Nanoscale 15, 14043–14054, doi:10.1039/d3nr01873c (2023).

69 Simmel, F. C. Nucleic acid strand displacement – from DNA nanotechnology to translational regulation. RNA Biology 20, 154–163, doi:10.1080/15476286.2023.2204565 (2023).

70 Ratajczyk, E. J., Šulc, P., Turberfield, A. J., Doye, J. P. K. & Louis, A. A. Coarse-grained modeling of DNA–RNA hybrids. The Journal of Chemical Physics 160, doi:10.1063/5.0199558 (2024).

71 Jung, J. K., Archuleta, C. M., Alam, K. K. & Lucks, J. B. Programming cell-free biosensors with DNA strand displacement circuits. Nature Chemical Biology 18, 385–393, doi:10.1038/s41589-021-00962-9 (2022).

72 Smith, F. G., Goertz, J. P., Jurinović, K., Stevens, M. M. & Ouldridge, T. E. Strong sequence–dependence in RNA/DNA hybrid strand displacement kinetics. Nanoscale 16, 17624–17637, doi:10.1039/d4nr00542b (2024).

73 Mills, A. et al. A modular spring-loaded actuator for mechanical activation of membrane proteins. Nat Commun 13, 3182, doi:10.1038/s41467-022-30745-2 (2022).

74 Ji, J., Karna, D. & Mao, H. DNA origami nano-mechanics. Chem Soc Rev 50, 11966–11978, doi:10.1039/d1cs00250c (2021).

75 Jeon, B.-j., et al. Modular DNA origami–based electrochemical detection of DNA and proteins. Proceedings of the National Academy of Sciences 122, doi:10.1073/pnas.2311279121 (2024).

76 Grabenhorst, L. et al. Engineering modular and tunable single-molecule sensors by decoupling sensing from signal output. Nature Nanotechnology 20, 303–310, doi:10.1038/s41565-024-01804-0 (2024).

77 Walbrun, A. et al. Single-molecule force spectroscopy of toehold-mediated strand displacement. Nature Communications 15, doi:10.1038/s41467-024-51813-9 (2024).

78 Platnich, C. M. et al. Kinetics of Strand Displacement and Hybridization on Wireframe DNA Nanostructures: Dissecting the Roles of Size, Morphology, and Rigidity. ACS Nano 12, 12836–12846, doi:10.1021/acsnano.8b08016 (2018).

79 Matin, T. R., Heath, G. R., Huysmans, G. H. M., Boudker, O. & Scheuring, S. Millisecond dynamics of an unlabeled amino acid transporter. Nature Communications 11, doi:10.1038/s41467-020-18811-z (2020).

80 Lorenz, R. et al. ViennaRNA Package 2.0. Algorithms for Molecular Biology 6, doi:10.1186/1748-7188-6-26 (2011).

81 Jazbutyte, V. & Thum, T. MicroRNA-21: From Cancer to Cardiovascular Disease. Current Drug Targets 11, 926–935, doi:10.2174/138945010791591403 (2010).

82 Letafati, A. et al. MicroRNA let-7 and viral infections: focus on mechanisms of action. Cellular & Molecular Biology Letters 27, doi:10.1186/s11658-022-00317-9 (2022).

83 Atif, H. & Hicks, S. D. A Review of MicroRNA Biomarkers in Traumatic Brain Injury. Journal of Experimental Neuroscience 13, doi:10.1177/1179069519832286 (2019).

84 Zou, R. et al. Development and validation of a circulating microRNA panel for the early detection of breast cancer. Br J Cancer 126, 472–481, doi:10.1038/s41416-021-01593-6 (2022).

85 Zhu, C. et al. A five-microRNA panel in plasma was identified as potential biomarker for early detection of gastric cancer. Br J Cancer 110, 2291–2299, doi:10.1038/bjc.2014.119 (2014).

86 Wang, C. et al. A Five-miRNA Panel Identified From a Multicentric Case-control Study Serves as a Novel Diagnostic Tool for Ethnically Diverse Non-small-cell Lung Cancer Patients. EBioMedicine 2, 1377–1385, doi:10.1016/j.ebiom.2015.07.034 (2015).

87 Ivanov, A. P. et al. On-Demand Delivery of Single DNA Molecules Using Nanopipets. ACS Nano 9, 3587–3595, doi:10.1021/acsnano.5b00911 (2015).

88 Kowalczyk, S. W., Wells, D. B., Aksimentiev, A. & Dekker, C. Slowing down DNA Translocation through a Nanopore in Lithium Chloride. Nano Letters 12, 1038–1044, doi:10.1021/nl204273h (2012).

89 Cruz-León, S. et al. Twisting DNA by salt. Nucleic Acids Research 50, 5726–5738, doi:10.1093/nar/gkac445 (2022).

90 Baumann, C. G., Smith, S. B., Bloomfield, V. A. & Bustamante, C. Ionic effects on the elasticity of single DNA molecules. Proceedings of the National Academy of Sciences 94, 6185–6190, doi:10.1073/pnas.94.12.6185 (1997).

91 Gallego Romero, I., Pai, A. A., Tung, J. & Gilad, Y. RNA-seq: impact of RNA degradation on transcript quantification. BMC Biology 12, doi:10.1186/1741-7007-12-42 (2014).

92 Chau, C., Mohanan, G., Macaulay, I., Actis, P. & Wälti, C. Automated Purification of DNA Origami with SPRI Beads. Small, doi:10.1002/smll.202308776 (2023).

93 Confederat, S. et al. Next-Generation Nanopore Sensors Based on Conductive Pulse Sensing for Enhanced Detection of Nanoparticles. Small, doi:10.1002/smll.202305186 (2023).

