## Supporting Information for "Visualizing and Quantifying microRNA Induced DNA Origami Separation at the Nanoscale"

### Method

Unless otherwise specified, all chemicals were obtained from Sigma-Aldrich at ACS grade.

#### Additional details on DNA origami

##### Left and Right Monomer

The design of the asymmetrical Tile DNA origami was based on the previously published 4-Fold Symmetrical Tile (4FST) by Tikhomirov *et al.* [2]. and 4-Fold Symmetrical Frame (4FSF) by Chau *et al.* [3]. Owing to their symmetries, the 4FST and 4FSF could both be broken down into modules of 4 triangles and 4 trapeziums respectively, we termed these modules as T1, T2, T3 and T4, each of these modules contained its own edge, interior and bridge staples mixture. To form the DNA origami monomers, the T1, T2 and T4 modules of the 4FST were mixed with the T3 modules of the 4FSF. The formation of the left monomer required a different set of edge staples for the T1 edge, similarly, the formation of the right monomer required a different set of edge staples for the T4 edge.

##### Linkers and combinations

The linkers sequences could be found as an excel file in the associated repository. All linker modifications happened at the edge of the origamis.

##### Dimer displacement

To displace the dimer with invaders, 2 nM (9.4 ng/μl by concentration) of dimer was mixed with various concentration of invaders, the invader sequence depended on the linker sequence. The mixture of invader and dimer would then be incubated at either 25°C or 37°C inside a thermal cycler for 30 minutes, then the temperature dropped to 4°C, the resultant samples were stored at 4°C.

##### Displaced dimer strand negation

The displacement of the dimer followed exactly as the above section. The strand negations were achieved by mixing a final concentration of 200 nM negation strand (a short complementary strand that hybridise to the exposed sticky end of the right monomer after the displacement), the mixtures were then incubated inside a thermal cycler for 30 minutes at 37°C, then the temperature dropped to 4°C.

##### RNAse digestion

The RNAse digestion was carried out by adding the RNAse Cocktail (AM2286; Thermo Fisher Scientific) or RNAse H (M0297S; NEB). The final concentration of the RNAse Cocktail contained 50U/ml of RNAse A and 2,000 U/ml of RNAse T1, the final concentration of the RNAse H contained 500 U/ml. The mixtures were incubated inside a thermal cycler for 30 minutes at 25°C, then the temperature dropped to 4°C.

### Absorption Spectroscopy and molarity conversion

The concentration of the dsDNA was measured via absorption spectroscopy using a NanoDrop™ 2000c Spectrophotometer (Thermo Fisher). The instrument was blanked with 1× TE buffer. 2 µl of the sample was dotted onto the measurement platform, and the measurement was performed using an extinction coefficient of 38 mg/ml for $A_{260}$ = 1. The concentration to molarity conversion was carried out by assuming molecule as ssDNA.

### Agarose Gel Electrophoresis

For the agarose gel analysis of the DNA origami, a 1.5% magnesium containing agarose gel was used. The high purity agarose (16500500; Thermo Fisher) was mixed with 0.5X TBE, 10 mM $MgCl_2$ buffer in an Erlenmeyer flask. The agarose was melted in microwave oven at full power for 45 seconds, and immediately poured into the casting module of the agarose gel electrophoresis system (Mini-Sub Cell GT Systems; Bio-Rad). The total loaded DNA mass was 25 ng per sample. The purified DNA origami was mixed with 2 µl of the loading dye (B7025S; NEB) and the volume was brought up to a total of 12 µl with DNA origami buffer. The M13mp18 ssDNA scaffold was prepared in the same way with a fixed mass of 50 ng for gel analysis. The gel was submerged in the 0.5X TBE, 10 mM $MgCl_2$ buffer and the samples were loaded with pipette, and 70 V was applied for 75 minutes. The gel was then transferred to a foil-covered container containing 30 ml of 1× TAE buffer, and 10 µl of the Diamond nucleic acid dye (H1181; Promega) was used to stain the gel for 2 hours under constant agitation prior imaging. The gel densitometry is processed by ImageJ.

### High-Speed AFM Imaging and Data Analysis

All high speed-atomic force microscopy (HS-AFM) measurements were performed using a NanoRacer HS-AFM (Bruker, Germany) instrument in amplitude modulation mode. All HS-AFM measurements were obtained in liquid and ambient temperature in an acoustic isolation housing on an active antivibration table using short cantilevers (USC-F1.2-k0.15, NanoWorld, Switzerland) with nominal spring constants of 0.15 $N.m^{-1}$, resonance frequencies of ~0.6 MHz and quality factors of ~2. To prepare the sample for HS-AFM imagining of DNA Origami, freshly cleaved mica was treated with 4 µl of 10 mM $Ni^{+}$/ $Mg^{2+}$ salt solution, to promote the adsorption of DNA origami to the mica. Thereafter, 2-4 µl of DNA Origami sample was incubated on the mica and left for incubation for 2-3 minutes before rinsing, with 5 via fluid exchange with 5 µl of buffer solution. The sample holder was then filled with 1 ml of imaging buffer (5 mM Tris-HCl (pH 8.0), 1 mM EDTA, 20 mM $MgCl_2$ and 5 mM NaCl). To visualize TMSD, 10 µl of the invader strands at a concentration of 1 µM were added to the imaging buffer. The images and videos were analysed with NanoLocz image analysis software (v1.30)[63].

HS-AFM data analysis including image leveling, alignment and Localization AFM of origami data and image/movie export was performed in NanoLocz (https://github.com/George-R-Heath/NanoLocz) AFM image analysis software [57] and thereafter image/movies were exported for further analysis. The obtained levelled and aligned movies were further imported in Fiji-imageJ software in Tiff format to generate the Kymograph profiles of the movies. To obtain a Kymograph profile, the origami dimer bridge connection was vertically aligned, and a vertical line was drawn on the linker connection, so that it tracks the movement of all 5 linker together and their separation profile over the time. The image reslice tool was then used to generate an orthogonal Kymograph profile. The generate grey scale kymograph profile shows the progress of each separation event over the linkers with time. Further a height profile was generated from this kymograph to easily visualize all the on/off states during separation process. Further, python script was used to define the on/off separation state with optimum single threshold value for all the linkers to demonstrate the separation in on/off states and to plot the final graphs (Figure 3c and Supporting Figures 15-19(d)). To add the AFM colors to grey scaled processed imaged, lookup table were used (https://github.com/George-R-Heath/AFM-LUTS).

### Nanopore population analysis

#### Kernel Density Estimation and Area under the curve

The Kernel Density Estimation (KDE) is a non-parametric statistical technique used to estimate the probability density function of a random variable from data [4]. The method involves placing a gaussian kernel at each data point and summing them to create a continuous curve that estimate the underlying distribution.

We utilised the KDE on the peak area of the translocation data to extrapolate the distribution of the data. The KDE bandwidth was determined by the Scott's rule of thumb method: $bw = (\frac{4\times\sigma^5}{3\times n})^{0.2}$ , where the bw was the bandwidth, $\sigma$ was the standard deviation of the data and n was the amount of data. Two bounds: the upper and the lower bounds were established by taking the peak value of the dimer population and the ±20% of that peak values were computed respectively. The area under the curve (AUC) of a probability density function represents the probabilities and sum to 1 (i.e. 100%). Through integration, the area under the curve (AUC) and the probabilities within bounds could be computed and thus the probability of a translocation event fell into the region that was classified as a dimer.

### Scanning Electron Microscopy

The nanopore dimensions were imaged by Scanning Electron Microscopy (SEM) using a Nova NanoSEM. The imaging was performed by Dr Alexander Kulak, University of Leeds.

### Fluorescence measurement and kinetic fitting

#### Measurement

The kinetic loop measurement was carried out in the Spark multimode microplate reader (TECAN) with excitation set at 485 nm and emission at 535 nm, the black bottom 96 wells plate (655076; Greiner) was used. A total of 8 wells were used for each measurement, and a total of 60 µl would be used for each well, and each well contained 2 nM (9.4 ng/µl) of dimer and invaders at different concentrations. All measurements were carried out at 25°C. One of the wells contained displaced fluorescent dimer , this sample was generated by adding the ssDNA invader at 100 nM and incubated for 1 hour before the measurement, the Z height and the gain optimisation would be carried out on this sample. One of the well contained the dimer without the addition of ssDNA invaders. After the plate was loaded into the microplate reader, it performed double orbital shakes for 5 seconds followed by calibration of the Z height and gain optimisation. 360 kinetic loops were performed, each loop lasted for 30 seconds, in each loop, each well was read 4 times at different points with 10 flashes each, after all the wells were read, 12 seconds of double orbital shaking would then be performed, followed by incubation time until the next loop to begin. The total read time was 3 hours for each kinetic loop measurement.

#### Fitting

The following equation was used to describe the TMSD kinetics carried out:

**Equation 1:**

$$Invader + Gate \xrightarrow{k_{eff}} Incumbent + Signal$$

Where Invader represented the ssDNA or the RNA invaders, the Gate represented the origami dimer, the Incumbent represented the left monomer where the invader hybridised with, and the Signal also represented the left monomer where the fluorophore (fluorescein) emitted fluorescent signal when excited with appropriate wavelength after the displacement had happened. Utilising the rate law, the following ordinary differential equations (ODEs) could be derived:

**Equation 2:**

$$\frac{dInvader}{dt} = -k_{eff} \times [Invader] \times [Gate]$$

**Equation 3:**

$$\frac{dGate}{dt} = -k_{eff} \times [Invader] \times [Gate]$$

**Equation 4:**

$$\frac{dIncumbent}{dt} = k_{eff} \times [Invader] \times [Gate]$$

**Equation 5:**

$$\frac{dSignal}{dt} = k_{eff} \times [Invader] \times [Gate]$$

The Equation 2-5 formed a set of ODEs, with python, we defined a function TMSD_ODEs_system. This function defined the above set of ODEs with three function inputs: time_t - array of time in seconds, Initial_concentration - initial Gate concentrations, $k_{eff}$ - rate constant. The function output is the derivatives: $\frac{dInvader}{dt}$, $\frac{dGate}{dt}$, $\frac{dIncumbent}{dt}$ and $\frac{dSignal}{dt}$, this function would then be solved by the scipy.integrate.odeint in python by using the TMSD_ODEs_system as the function input. The scipy.optimize.curvefit took the integral of $\frac{dSignal}{dt}$, time_t and the initial concentration of Gate as inputs and utilised the non-linear least squares approach to fit the integral of the $\frac{dSignal}{dt}$ to the experimental data and estimated for $k_{eff}$ and α, α is a scaling factor. We provided an initial estimate of $k_{eff}$ and α values of 5 and 1, respectively. We provided upper and lower bounds for the fitted rate constants to be between 1 to 10 and the α between 0.005 to 1.0. The python code used to perform the fitting was modified based on Liu *et al.* [5] and could be accessed at https://github.com/chalmers4c/biochem_kinetic_fitting.

### RNA Minimum Free Energy prediction

The RNA minimum free energy prediction and its associated predicted secondary structures were computed through the ViennaRNA package [6] at http://rna.tbi.univie.ac.at. The predictions were carried out without folding constraints, avoid isolated base pairs, dangling energies on both sides of a helix in any case, using the Turner model, 2004 RNA parameters and temperature was set to either 25°C or 37°C at 0.01 M NaCl.

### Human tissue, tissue culture and RNA extraction

The human brain and kidney tissues were acquired through Thermo Fisher Scientific catalogue QS0611 and QS0616, respectively.

For the culture of HeLa cells, the ECACC authenticated HeLa cells (CVCL_0030) were cultured with Dulbecco's Modified Eagle's Medium (D5671) supplemented with 1X GlutaMax (35050038;

Thermo Fisher), 1X Penicillin Streptomycin (15140122; Fisher Scientific) and 10% (v/v) foetal bovine serum (F7524; Sigma Aldrich) inside cell culture flask (83.3911.302; Sarstedt) inside an incubator at 37°C with 5% $CO_2$. The cells were passaged when they were confluent, the cells were first washed with Magnesium, calcium free 1× DPBS (D8537), followed by the addition of a pre-warmed 1× trypsin-EDTA and left inside the incubator for 3 minutes, the detached cells were collected by centrifugation at 500×g to pellet the cells, the pellet was resuspended in medium then re-plated into a new flask. To cryopreserve the cells for long term storage, after the removal of the trypsin-EDTA solution, cells were resuspended in Synth-a-Freeze™ Cyropreservation medium (A1254201; Gibco) at minimum of $1X10^6$ cells/ml, and 1 ml of cell mixture was dispensed into a 1.8 ml cryovial (E3090-6222; Starlab). The cryovial was immediately sealed inside a polystyrene box and frozen down overnight at -80°C, and then transferred to a liquid nitrogen canister for long term storage. The extraction of the RNA was achieved with a commercially available extraction kit with the steps indicated in the kit (ReliaPrep™ miRNA Cell and Tissue Miniprep System; Z6210; Promega). For RNA extraction, all equipment and reagents were RNAse decontaminated or at RNAse-free grade.

### Supporting Figure

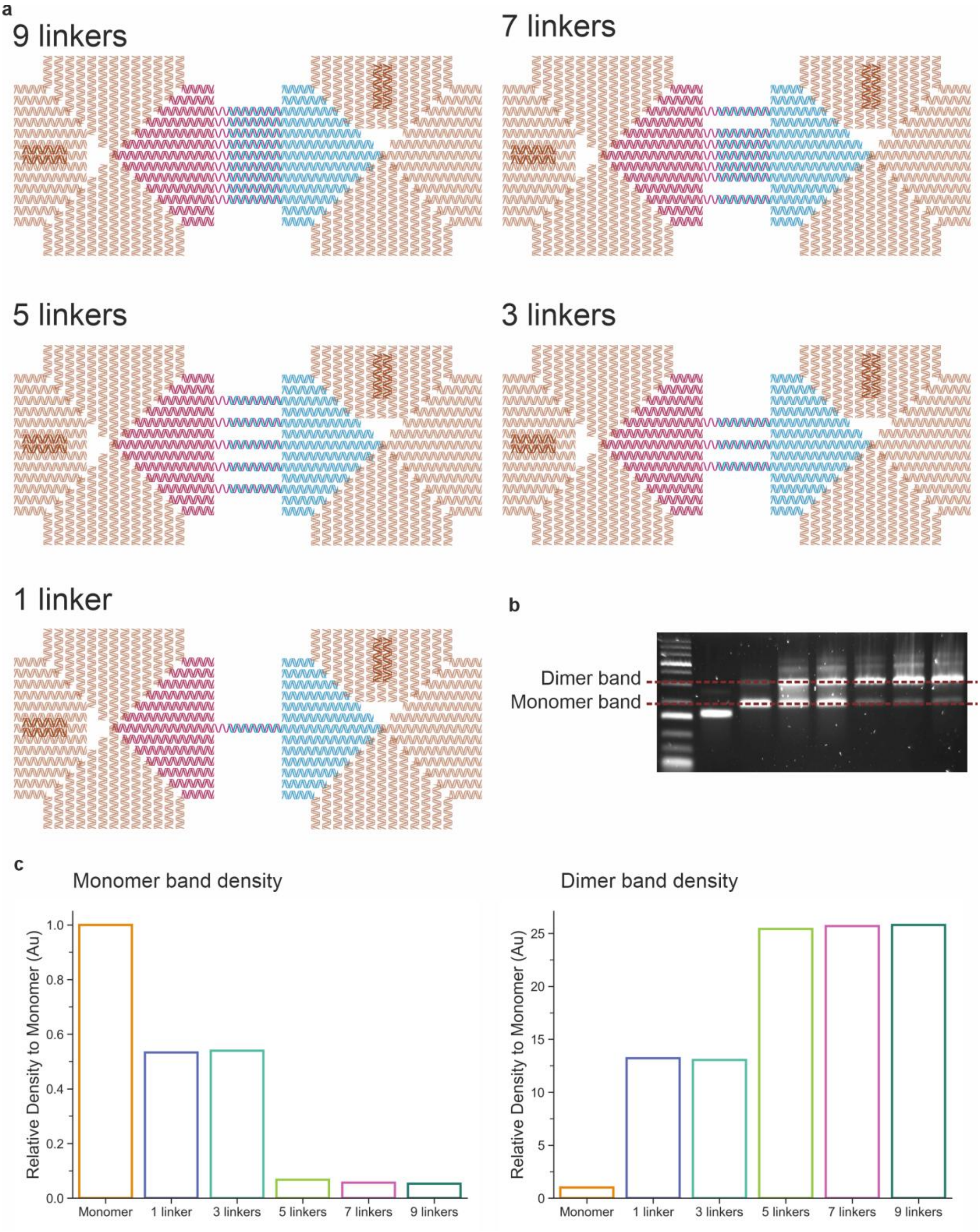

**Figure S1. Tested combinations of the DNA origami dimer and gel quantification.** (**a**) Schematic illustration of the linkers' positions and the number of linkers. (**b**) Position of the monomer band and dimer band. (**c**) Gel band densitometry quantification. The mass of total DNA loaded into the gel was constant across all samples (25 ng each lane), the monomer lane is used as the reference to calculate the relative densitometry, thus monomer has a relative density of 1 in both monomer band and dimer band density. For 5, 7 and 9 linkers, there are near 0 relative density at the monomer band position and near 25 times higher than monomer in the dimer band position. For 1 and 3 linkers, they have more monomers than 5, 7, 9 linkers and less dimer than 5, 7, 9 linkers.

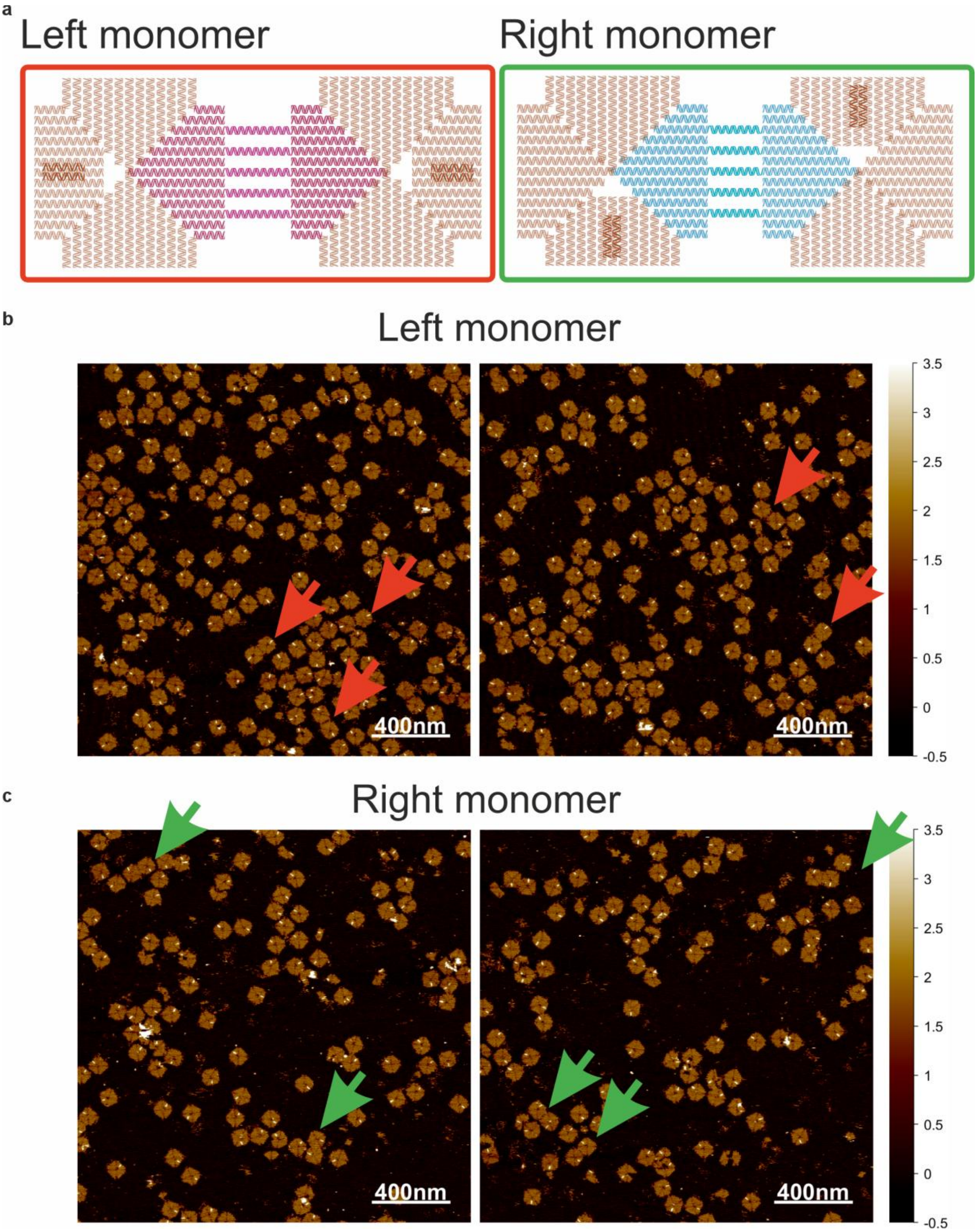

**Figure S2. The self-dimerization DNA origami dimer identified through AFM.** (**a**) schematic representations of the possible self-dimerization between the left and right monomers. Self-dimerized left monomer will have the orientation markers on the opposite end, along the longer axis. Self-dimerized right monomer will have the orientation markers on opposite end, along the short axis. (**b-c**) The AFM images of the left monomer and right showing the presence of self-dimerized dimers (arrows), respectively.

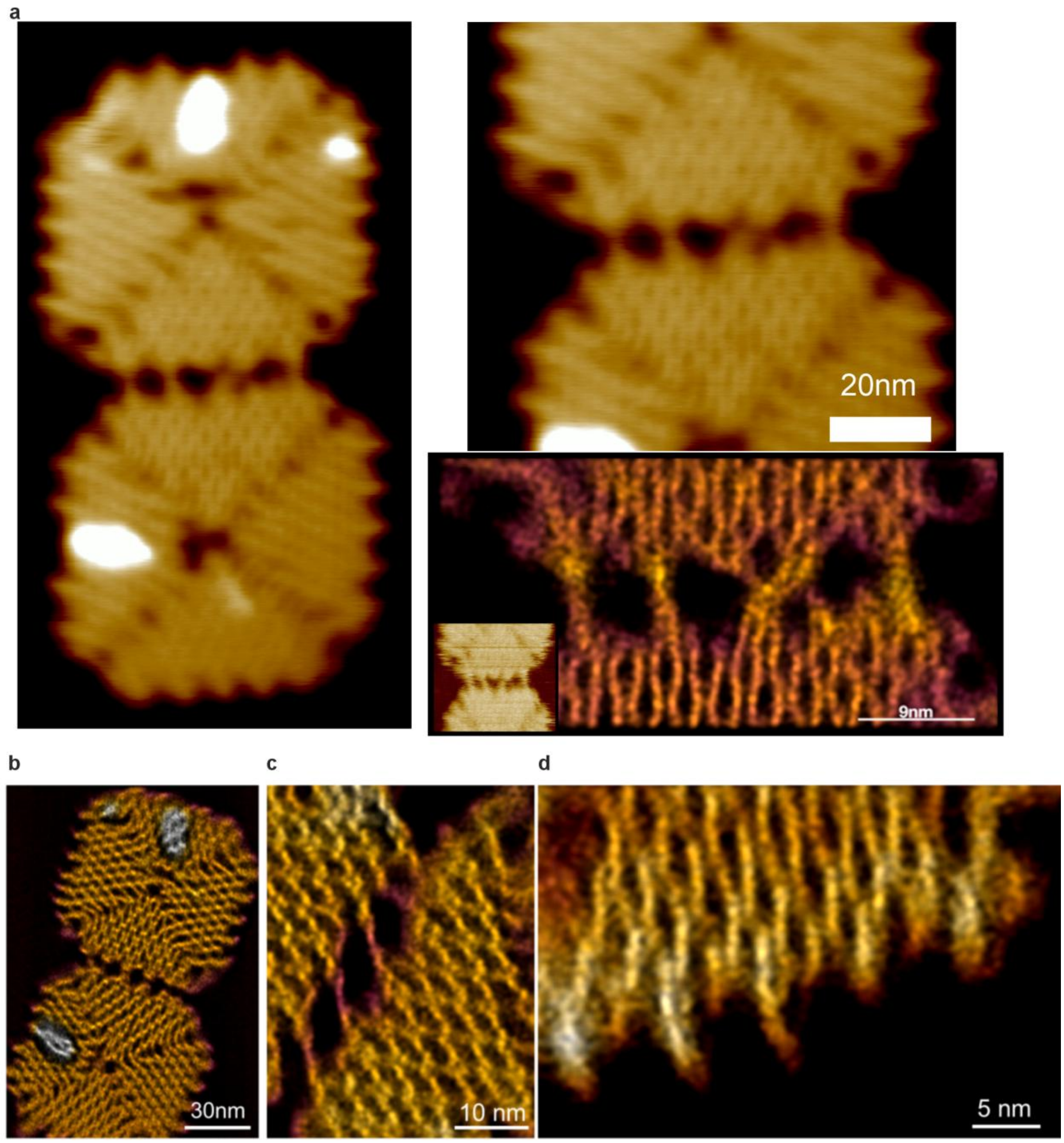

**Figure S3. Localization AFM images of origami dimers**. (**a**)At rare occasion, the linkers cross to form an "X" shaped connection between the two origami tiles. (**b**) LAFM image of the part the origami to visualize DNA strands folding. (**c**) L-AFM image of the dimer bridge, to visualize each of 5 linkers with higher resolution. (**d**) LAFM image of the origami, after successful TMSD process, to show the remaining linker strands on the origami.

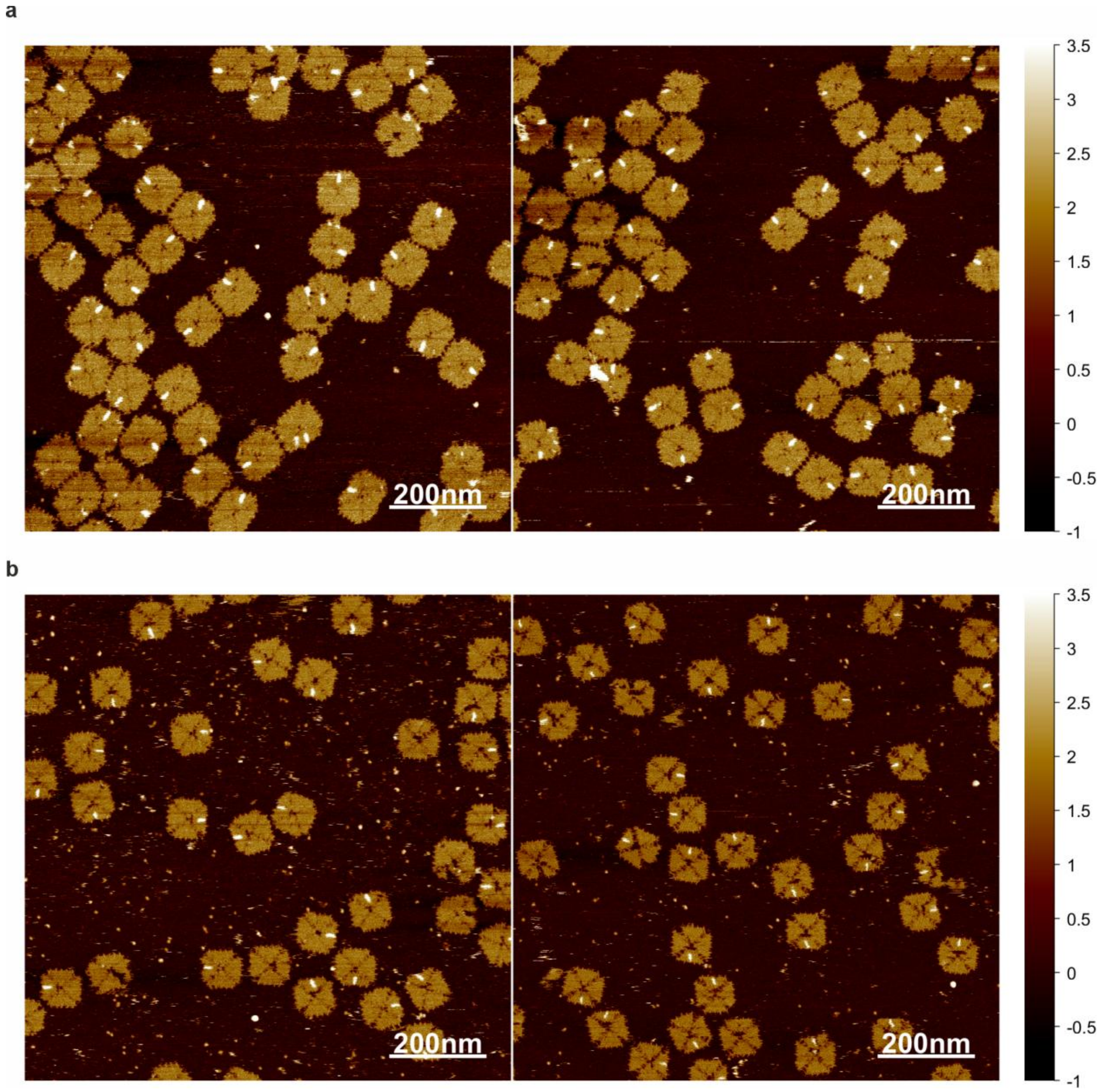

**Figure S4. Additional AFM images for the dimer and the displaced dimer.** (**a**) The dimer. (**b**) displaced dimer.

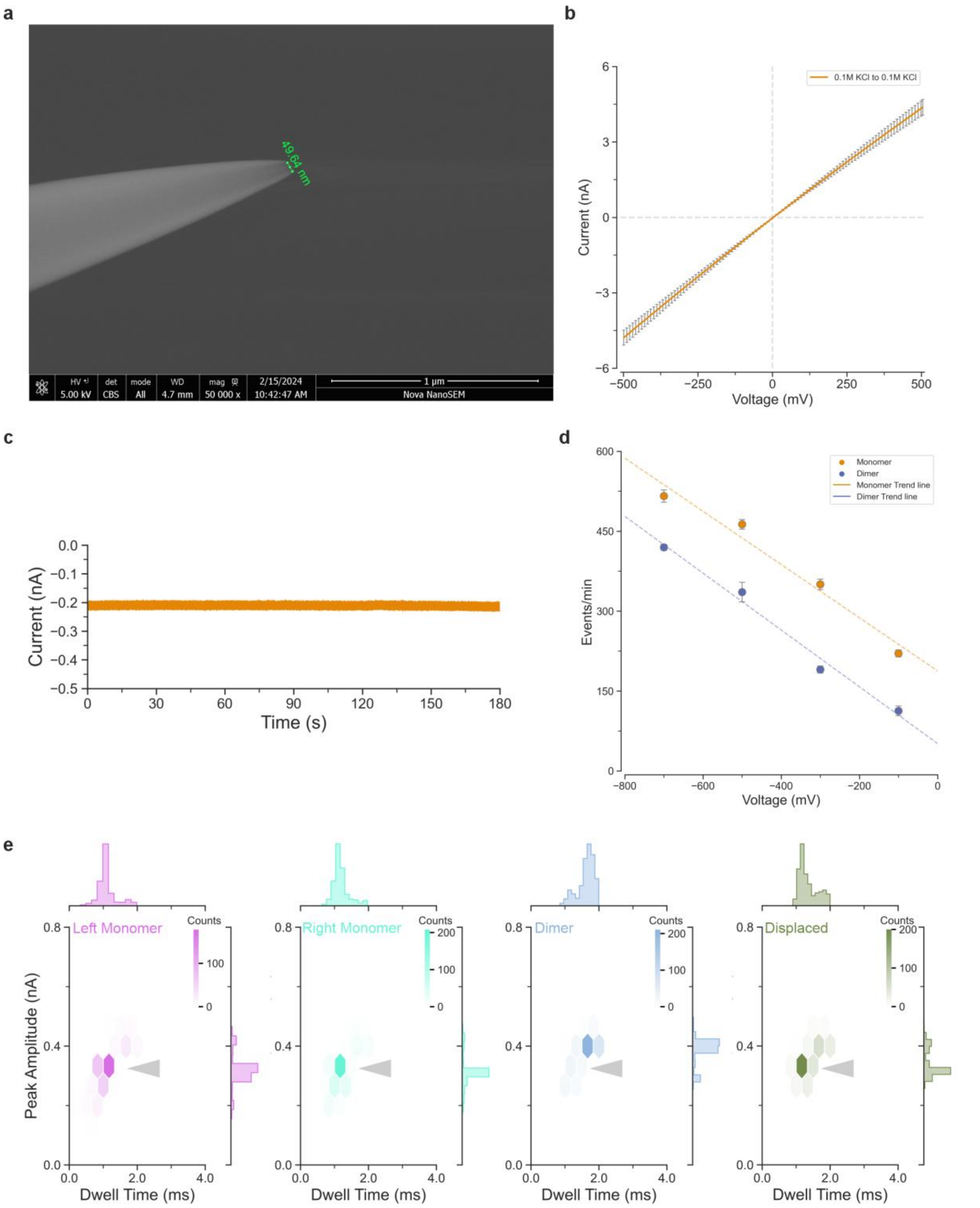

**Figure S5. Characterisation of the nanopore and translocations of DNA origamis.** (**a**)The micrograph shows a nanopore of approximately 50 nm in diameter. (**b**) The voltammetry measurements of 6 glass nanopores, both cis and trans chambers were filled with 0.1M KCl, the voltammetry shows that the nanopores are highly reproducible. Error bars are standard deviation. (**c**) Recording of the translocation trace when the cis chamber contains no analyte at -500 mV. (**d**) Translocation event rate for monomer and dimer across different voltages (n = 3, error bars are S.D.). (**e**) The population density plot for the detected translocation events. The population of the dimer shifted upward and outward comparing to the monomers, indicating increasing magnitude of current and event dwell time. The left monomer had a median of 1.1 ms for dwell time and 0.31 nA for current amplitude, the right monomer had a median of 1.2 ms for dwell time and 0.31 nA for current amplitude, the dimer had a median of 1.7 ms for dwell time and 0.4 nA for current amplitude, the displaced dimer had a median of 1.3 ms for dwell time and 0.32 nA for current amplitude.

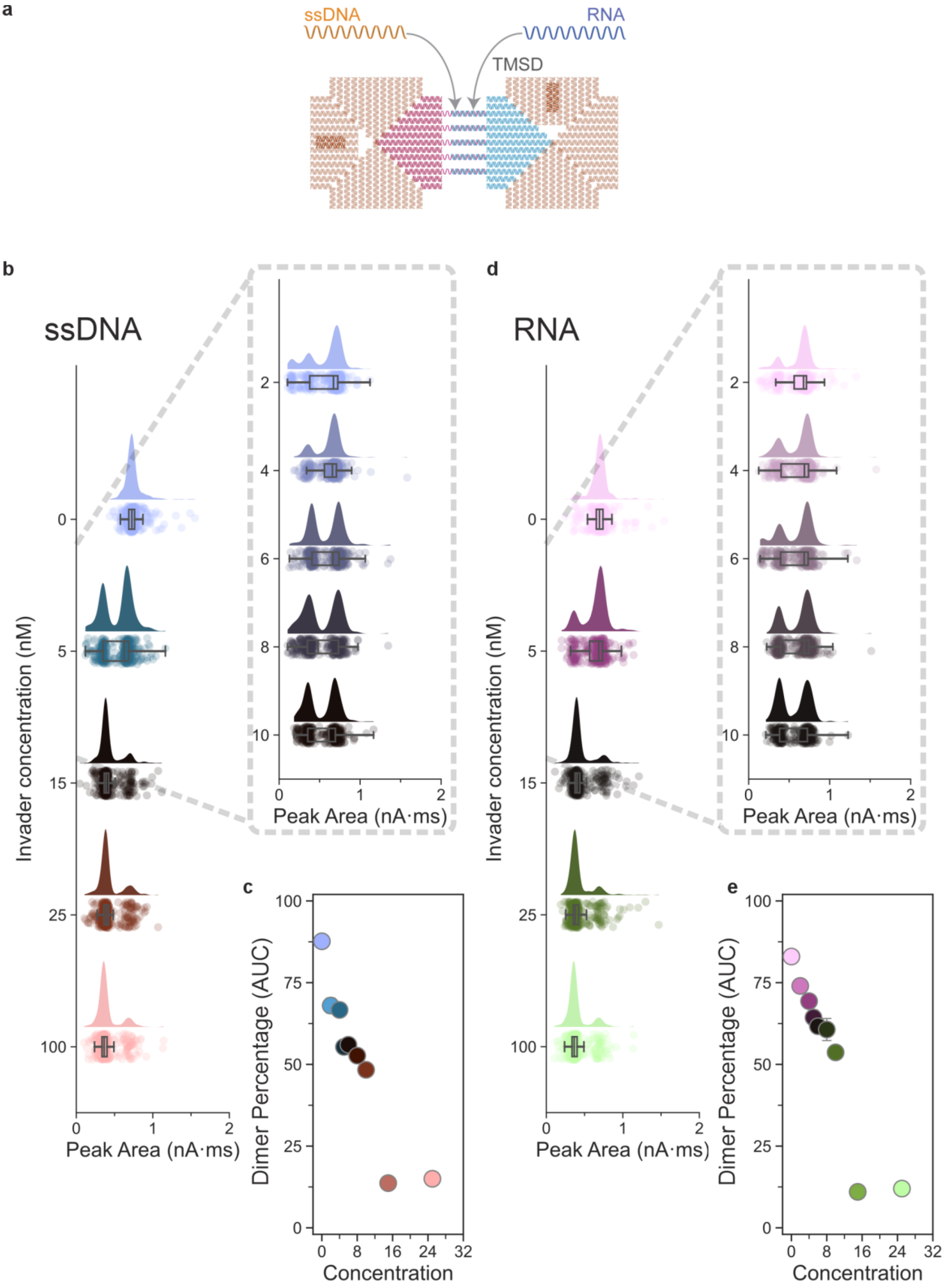

**Figure S6. The displacement caused by either ssDNA or RNA.** (**a**) The dimer were displaced by different concentration of invaders, the invaders were either the ssDNA (**b, c**) or the RNA (**d, e**), beside structural differences between ssDNA and RNA, the RNA invader had the same sequence as the ssDNA invader except the replacement of thymine with uracil. The ssDNA invader showed to be more effective at displacing the dimer as early on as 5 nM, the peak area shifted from approximately 0.7 nA·ms to approximately 0.4 nA·ms.

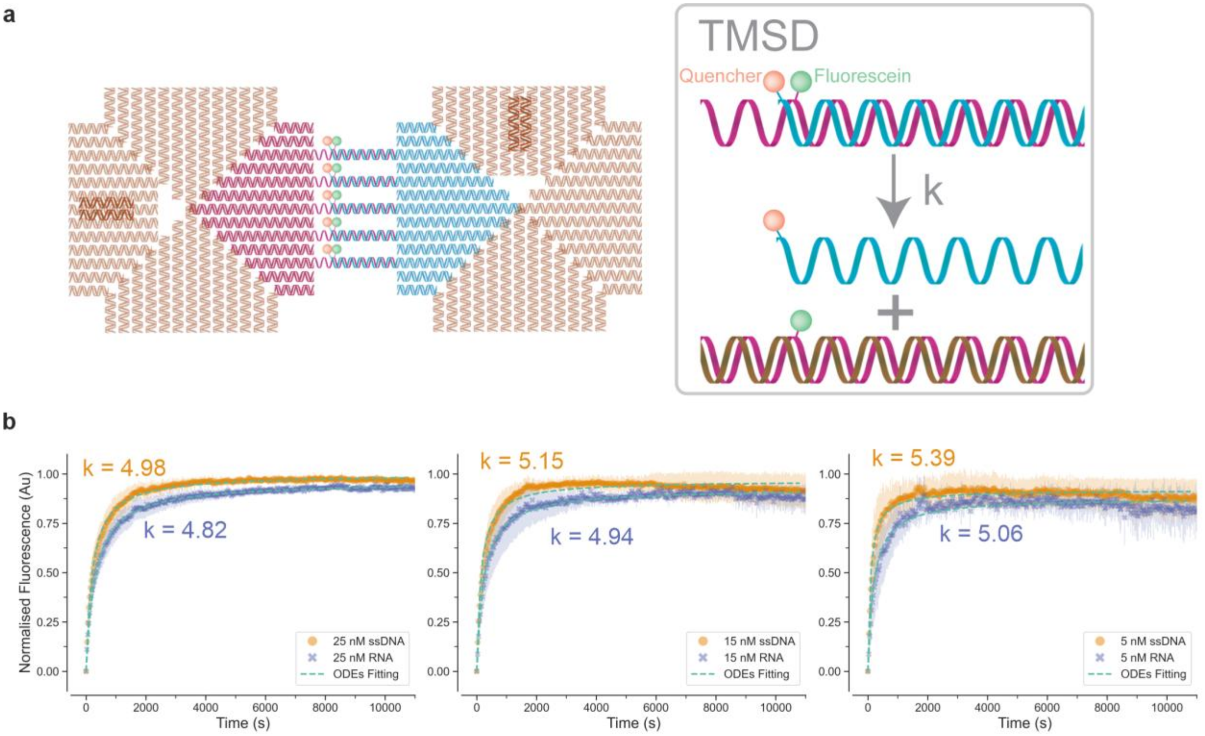

**Figure S7. TMSD kinetics.** (**a**) A specialised fluorophore-quencher dimer was synthesised, the right monomer had 5 quenchers (Iowa Black ® FQ) at the 5' position of the 5 linkers, and 5 fluoresceins were internally attached to the left monomer linkers such at that the quenchers were at close proximity with the fluoresceins upon dimerization. The parameter k is the rate constant from the displacement reaction. (**b**) The kinetics of the TMSD reactions within the dimer were monitored through a quencher and fluorophore (fluorescein) pair. The TMSD caused the displacement of the dimer , separating the quencher and fluorophore pair, the excitation of the fluorophore caused it to emit energy at higher wavelength. Both ssDNA and RNA invaders at 25, 15 and 5 nM reached equilibrium by 10800 seconds (3 hours), the rate constant for both reactions were fitted and computed through solving ordinary differential equations via integration, the ssDNA had a higher rate constant (k = 4.98 at 25 nM; k = 5.15 at 15 nM; k = 5.39 at 5 nM) while RNA had a slightly slower rate constant (k = 4.82 at 25 nM; k = 4.94 at 15 nM; k = 5.06 at 5 nM), the shaded region represented standard deviation.

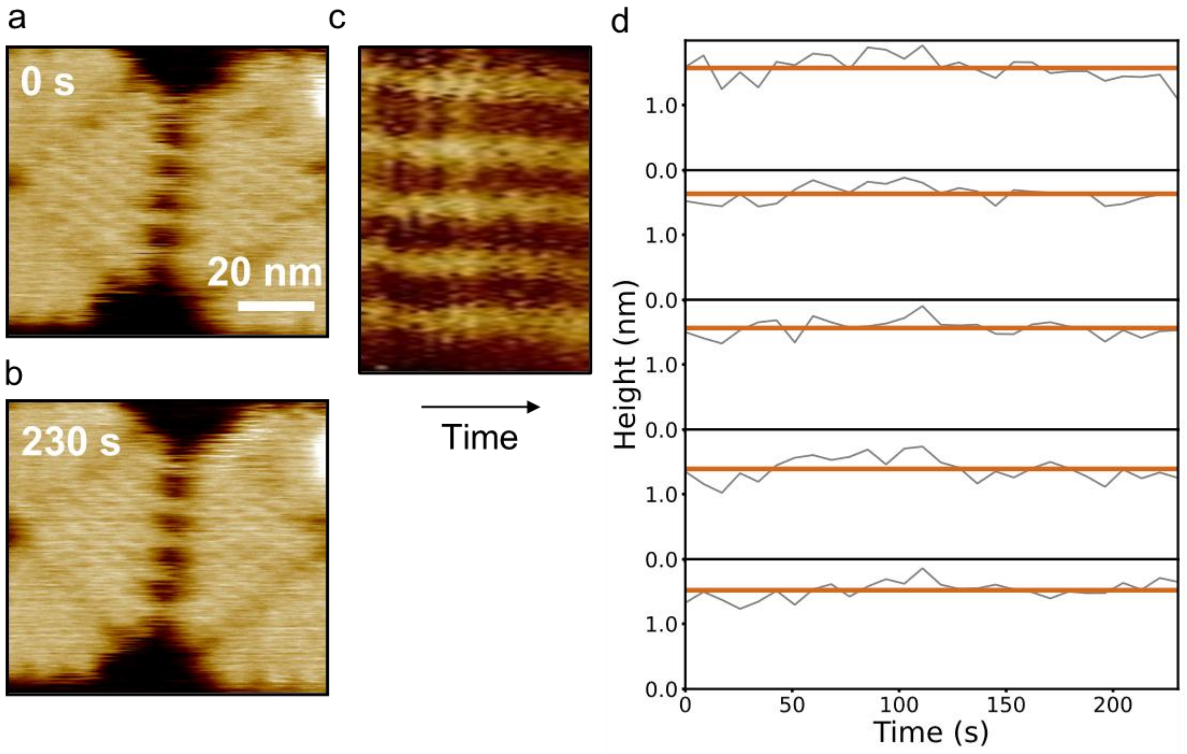

**Figure S8. High-speed imaging of the DNA origami, before addition of the DNA invader strands, as a control experiment.** HS-AFM image of the start (**a**) and end frame (**b**) of origami to confirm the stability of dimer during measurement. (**c**) Kymograph profile of all the linkers overtime, from the imaged movie (Supporting Movie 2). Height profile of the all the 5 linkers to show their single state confirmation.

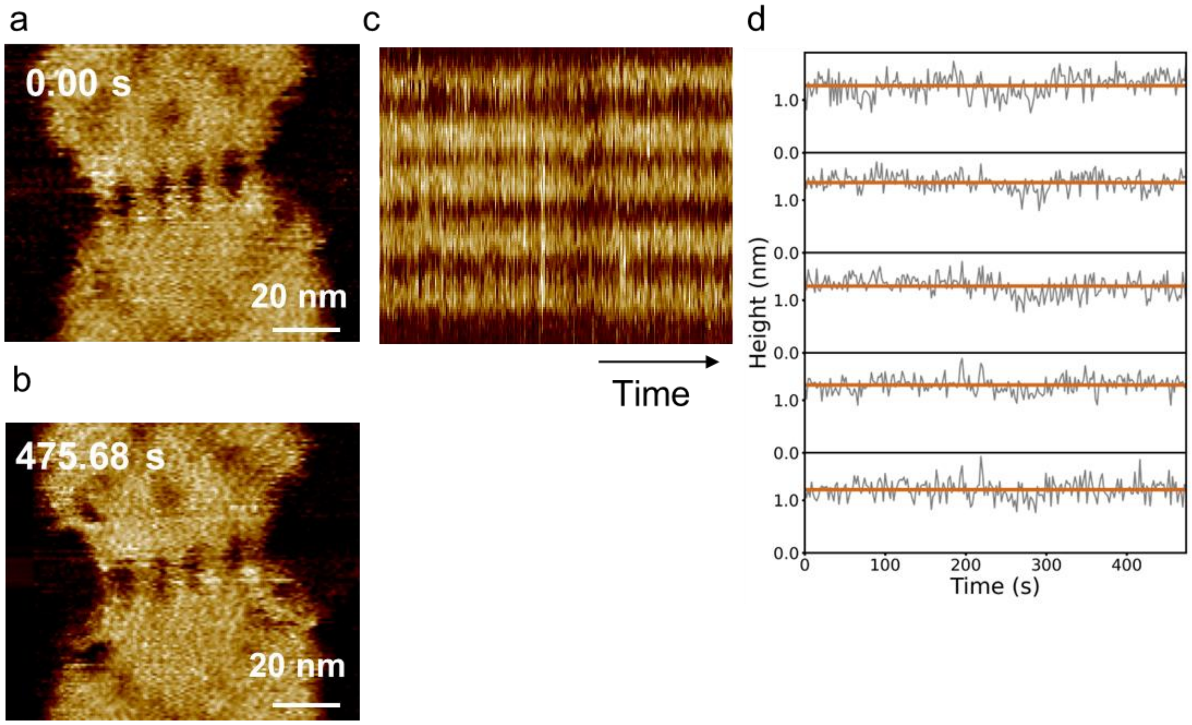

**Figure S9. High-speed imaging of the DNA origami, before addition of the RNA invader strands, as a control experiment.** HS- AFM image of the start (**a**) and end frame (**b**) of origami to confirm the stability of dimer during measurement. (**c**) Kymograph profile of all the linkers overtime, from the imaged movie (Supporting Movie 3). Height profile of the all the 5 linkers to show their single state confirmation.

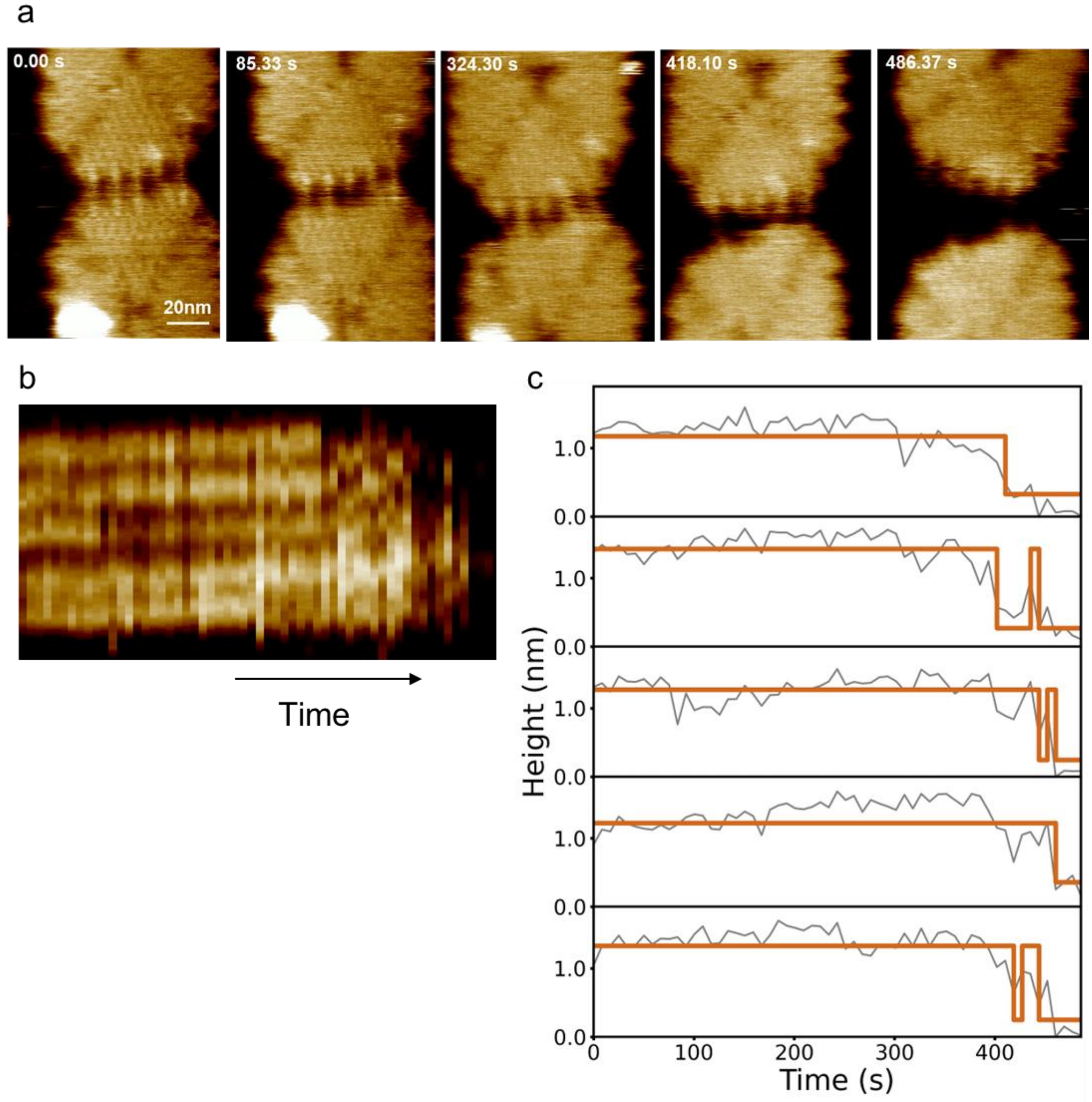

**Figure S10. High-speed imaging of the DNA origami after adding the DNA invader strands.** HS- AFM image of the start (**a**) Series of HS-AFM frames validating the successful TMSD separation of dimer (b) Kymograph profile of all the linkers overtime till successful TMSD separation, (Supporting Movie 5). (c) Height profile of the all the 5 linkers to show their on/off states overtime, till successful TMSD point.

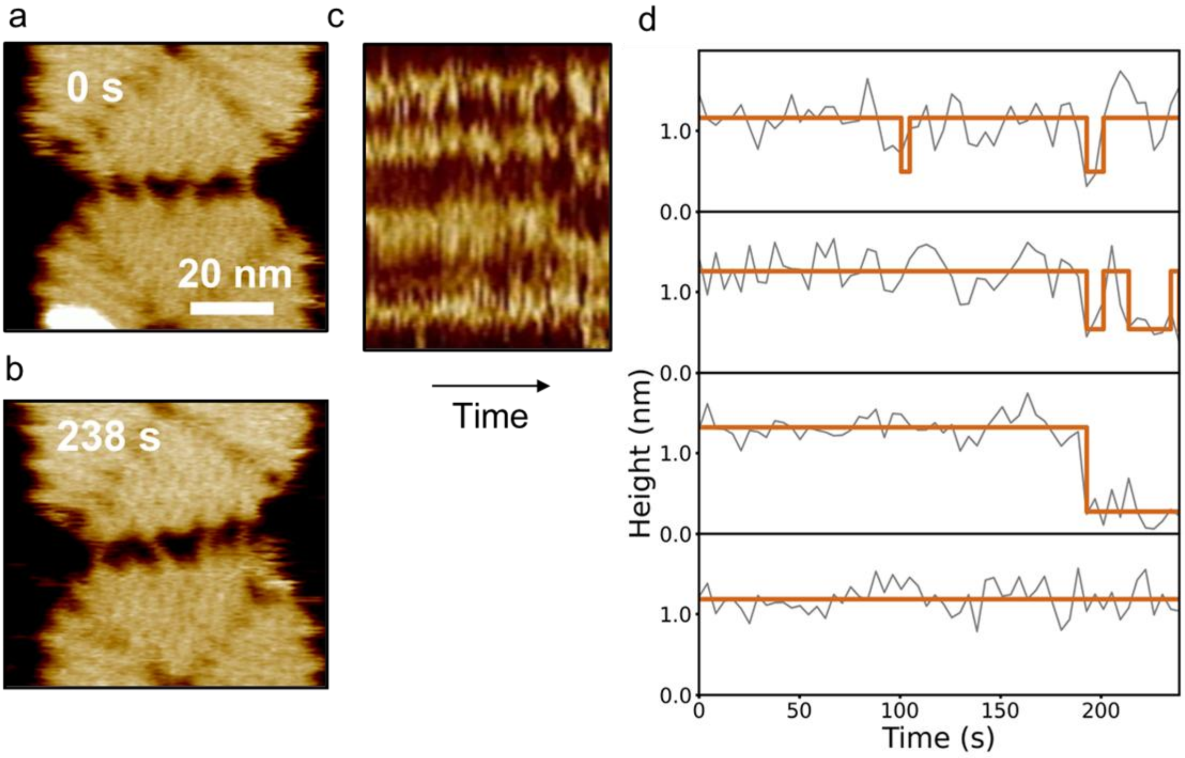

**Figure S11. High-speed imaging of the DNA origami after adding the DNA invader strands.** HS-AFM image of the start (**a**) and end frame (**b**) of origami to demonstrate the rare crossover event during partial TMSD process. (**c**) Kymograph profile of all the linkers overtime, from the imaged movie (Supporting Movie 6). Height profile of the all the 5 linkers to show their on/off states overtime.

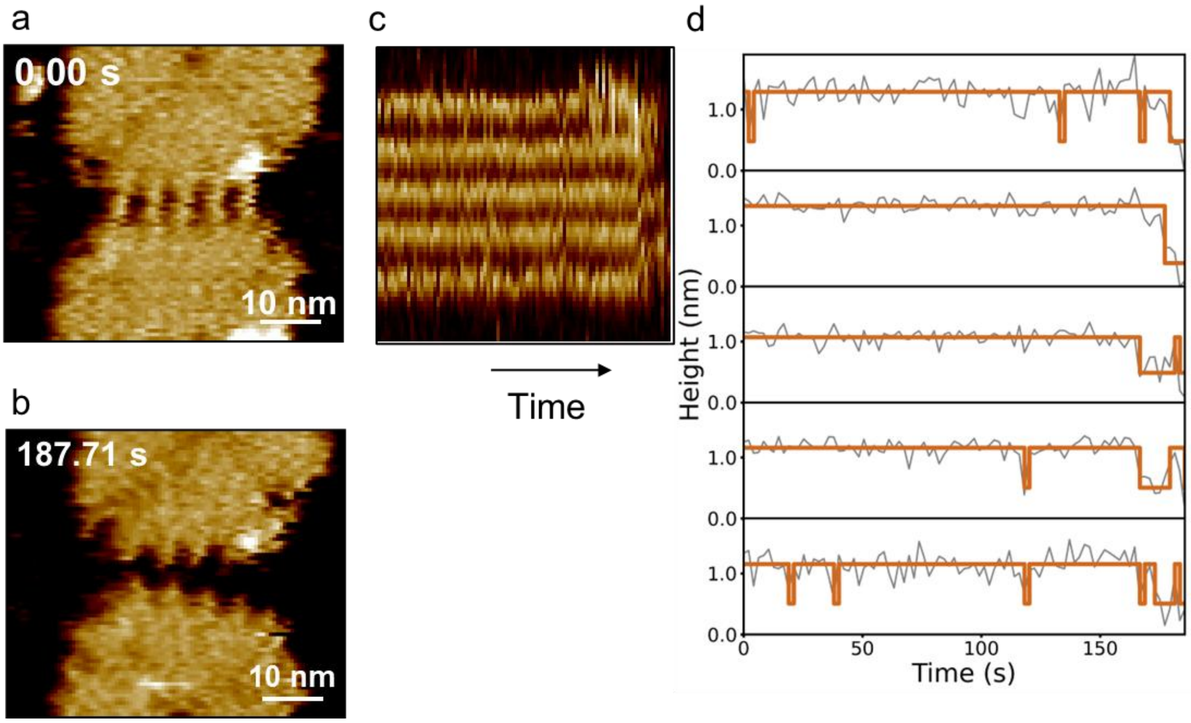

**Figure S12. High-speed imaging of the DNA origami after adding the RNA invader strands.** HS-AFM image of the start (**a**) and end frame (**b**) of origami, to demonstrate the successful TMSD process. (**c**) Kymograph profile of all the linkers overtime, from the imaged movie (Supporting Movie 7). Height profile of the all the 5 linkers to show their on/off states overtime.

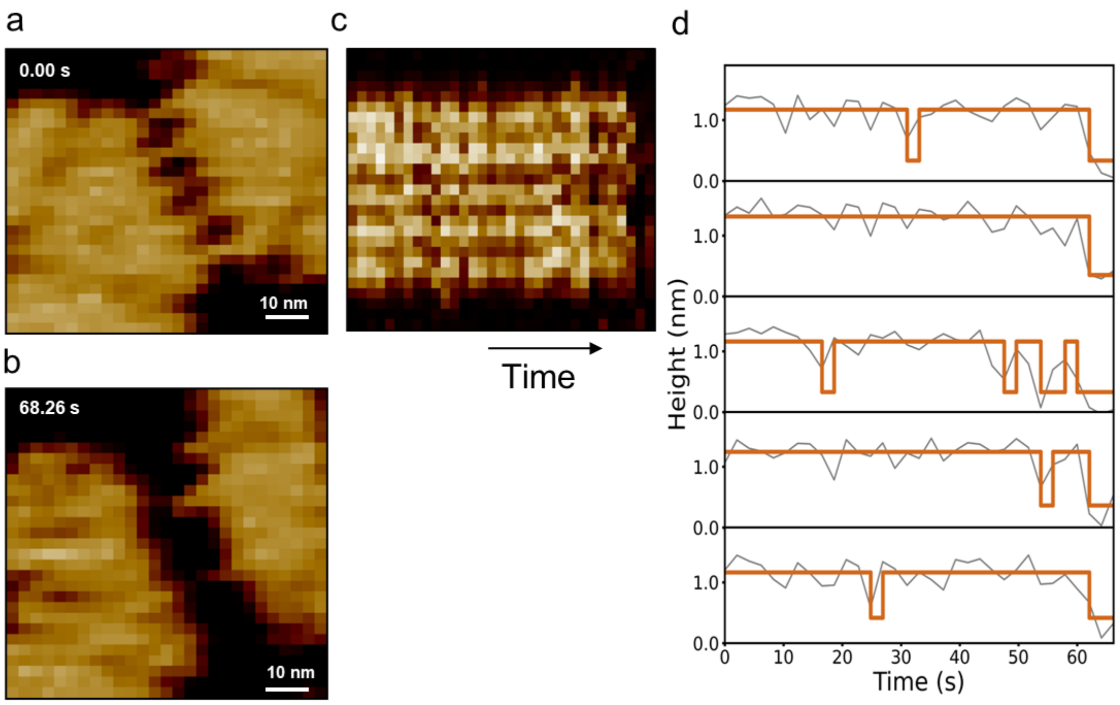

**Figure S13. High-speed imaging of the DNA origami after adding the RNA invader strands.** HS-AFM image of the start (**a**) and end frame (**b**) of origami to demonstrate the successful TMSD process. (**c**) Kymograph profile of all the linkers overtime, from the imaged movie (Supporting Movie 8). Height profile of the all the 5 linkers to show their on/off states overtime.

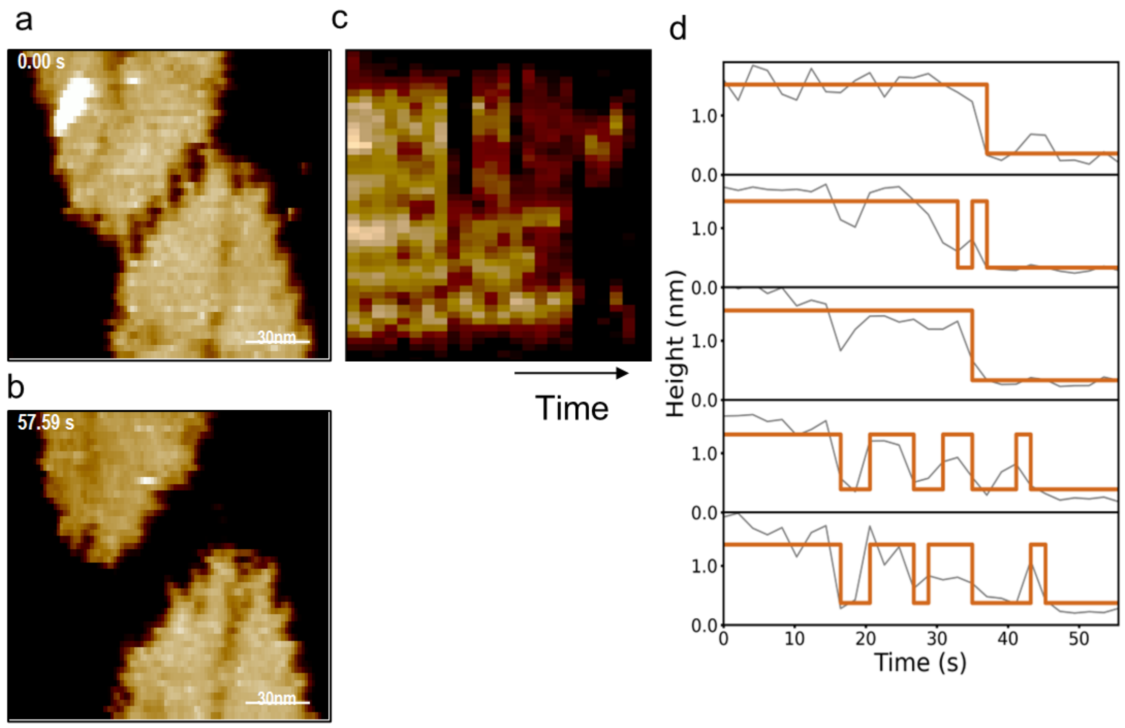

**Figure S14. High-speed imaging of the DNA origami after adding the RNA invader strands.** HS-AFM image of the start (**a**) and end frame (**b**) of origami, to demonstrate the successful TMSD process. (**c**) Kymograph profile of all the linkers overtime, from the imaged movie (Supporting Movie 9). Height profile of the all the 5 linkers to show their on/off states overtime.

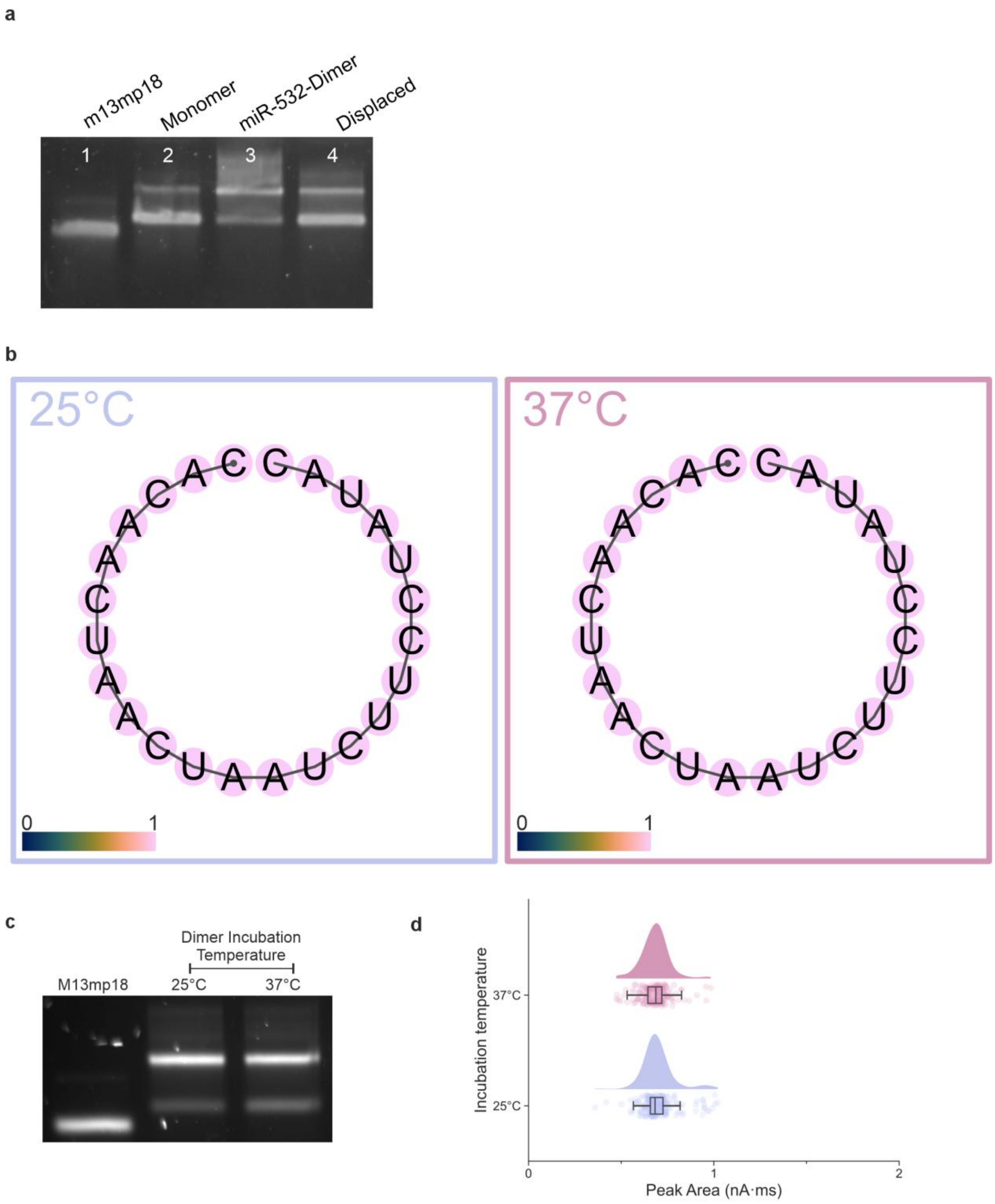

**Figure S15. Construction of the miR-532 dimer.** (**a**) Gel electrophoresis confirmed the construction of dimer with miR-532-5p linker sequence. The dimer underwent displacement when the invader strands (miR-532) were added. (**b**) The minimum free energy (MFE) structures of the SLD RNA invader at 10mM $Na^{+}$ conditions. The SLD RNA invader MFE structures showed that it does not form stable secondary structures. The colour gradient represents the base pairing probabilities (0 to 1), 0 being low pairing probabilities and 1 being high pairing probabilities, for unpaired regions, the colour denotes the probabilities of being unpaired with 1 being high probabilities of unpaired [6]. (**c**) Gel electrophoresis on the stability of the dimer when incubated at 25°C or 37°C for 30 minutes. (**d**) The calculated peak area of the dimer after incubating at either 25˚C or 37˚C, both samples have peak area of approximately 0.7 nA·ms. Indicating no significant degradation of the nanostructure after incubation.

**Supporting Table 1. ViennaRNA calculated MFE for miR-532-5p.**

| Temperature (˚C) | dot-bracket notation | MFE (kcal/mol) |
|---|---|---|
| 25 | ....(((.......)))..... | -1.06 |
| 26 | ....(((.......)))..... | -0.96 |
| 27 | ....(((.......)))..... | -0.87 |
| 28 | ....(((.......)))..... | -0.78 |
| 29 | ....(((.......)))..... | -0.7 |
| 30 | ....(((.......)))..... | -0.58 |
| 31 | ....(((.......)))..... | -0.51 |
| 32 | ....(((.......)))..... | -0.41 |
| 33 | ....(((.......)))..... | -0.33 |
| 34 | ....(((.......)))..... | -0.23 |
| 35 | ....(((.......)))..... | -0.14 |
| 36 | ....(((.......)))..... | -0.05 |
| 37 | ...................... | 0 |
| 38 | ...................... | 0 |
| 39 | ...................... | 0 |
| 40 | ...................... | 0 |

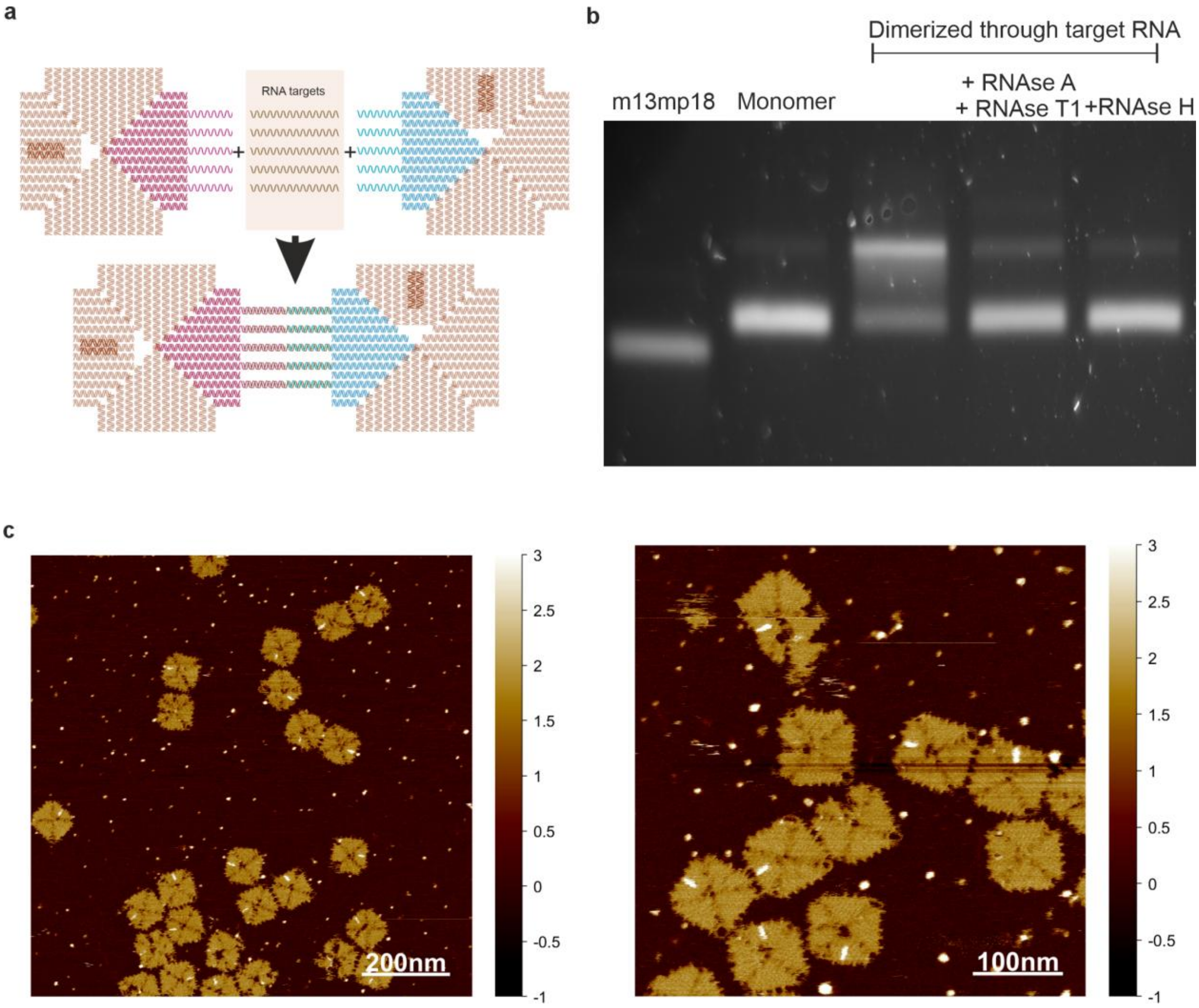

**Figure S16. The bottom-up construction approach.** (**a**) The DNA origamis can be dimerized by modifying the origami's edges sequence to become complementary to a target RNA where the left origami is complementary to half of the target RNA sequence and the right origami is complementary to the remaining half of the target sequence. Mixing the left monomer, target RNA and right monomer formed the dimer, since the structures transition from lower order to higher order, we termed this the bottom-up approach. (**b**) Agarose gel analysis on the stability of the bottom-up approach under the exposure of RNAse. The initially dimerized origamis returned to monomeric state, due to the linking target RNA was exposed to and subsequently digested by RNAse A, T1 and H at 25°C for 30 minutes. (**c**) AFM images of the dimer connected by a target RNA sequence using the approach in (a).

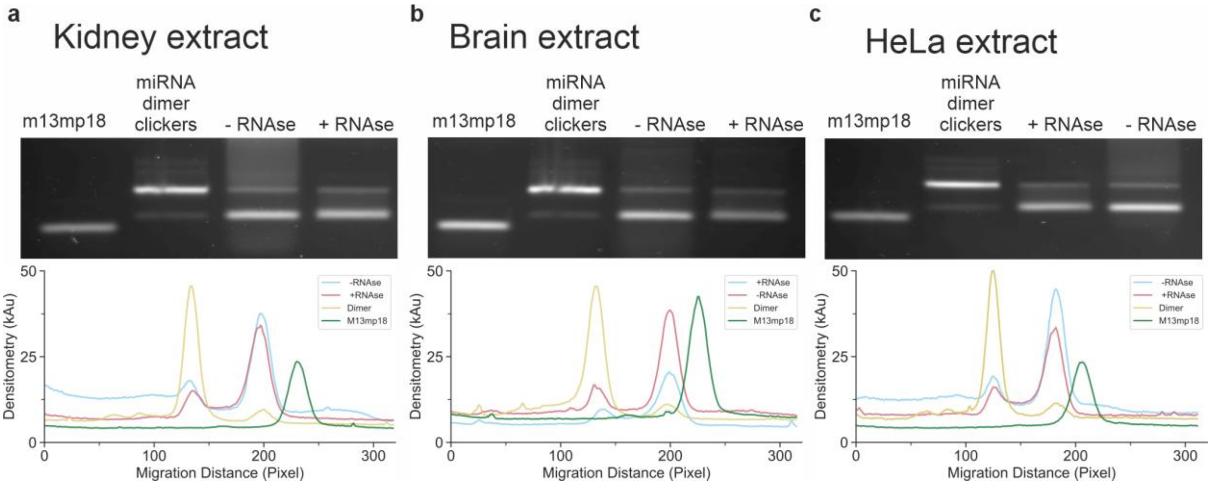

**Figure S17. The displacement of the dimer with RNA background and treatment of RNAse.** The miRNA dimer was exposed first to 5 nM of invader to cause TMSD at 37°C, followed by negation procedure at 37°C, then followed by RNAse digestion (or without RNAse digestion) at 37°C. In all cases (**a-c**) a noticeable background could be found in as a smear on the gel, these are the extracted RNA. The RNAse completely digested the RNA background. The lane profile was traced to visualise the higher densitometry background found in the lane without RNAse digestion. The RNAse digestion did not cause the dimer to be re-formed after the addition of the negation strand.

### References

1 Douglas, S. M. *et al.* Rapid prototyping of 3D DNA-origami shapes with caDNAno. *Nucleic Acids Research* **37**, 5001-5006, doi:10.1093/nar/gkp436 (2009).

2 Tikhomirov, G., Petersen, P. & Qian, L. Programmable disorder in random DNA tilings. *Nature Nanotechnology* **12**, 251-259, doi:10.1038/nnano.2016.256 (2016).

3 Chau, C., Mohanan, G., Macaulay, I., Actis, P. & Wälti, C. Automated Purification of DNA Origami with SPRI Beads. *Small*, doi:10.1002/smll.202308776 (2023).

4 Zielinski, W., Węglarczyk, S., Kuchar, L., Michalski, A. & Kazmierczak, B. Kernel density estimation and its application. *ITM Web of Conferences* **23**, 00037, doi:10.1051/itmconf/20182300037 (2018).

5 Liu, H. *et al.* Kinetics of RNA and RNA:DNA Hybrid Strand Displacement. *ACS Synthetic Biology* **10**, 3066-3073, doi:10.1021/acssynbio.1c00336 (2021).

6 Lorenz, R. *et al.* ViennaRNA Package 2.0. *Algorithms for Molecular Biology* **6**, doi:10.1186/1748-7188-6-26 (2011).

7 Charron, M., Briggs, K., King, S., Waugh, M. & Tabard-Cossa, V. Precise DNA Concentration Measurements with Nanopores by Controlled Counting. *Analytical Chemistry* **91**, 12228-12237, doi:10.1021/acs.analchem.9b01900 (2019).
